# Increased ACTL6A Occupancy Within mSWI/SNF Chromatin Remodelers Drives Human Squamous Cell Carcinoma

**DOI:** 10.1101/2021.03.22.435873

**Authors:** Chiung-Ying Chang, Zohar Shipony, Ann Kuo, Kyle M. Loh, William J. Greenleaf, Gerald R. Crabtree

## Abstract

Mammalian SWI/SNF (BAF) chromatin remodelers play dosage-sensitive roles in many human malignancies and neurologic disorders. The gene encoding the BAF subunit, ACTL6A, is amplified at an early stage in the development of squamous cell carcinomas (SCCs), but its oncogenic role remains unclear. Here we demonstrate that ACTL6A overexpression leads to its stoichiometric assembly into BAF complexes and drives its interaction and engagement with specific regulatory regions in the genome. In normal epithelial cells, ACTL6A was sub-stoichiometric to other BAF subunits. However, increased ACTL6A levels by ectopic expression or in SCC cells led to near-saturation of ACTL6A within BAF complexes. Increased ACTL6A occupancy enhanced polycomb opposition over the genome activating SCC genes and also enhanced the recruitment of transcription factor TEAD with its co-activator YAP, promoting their chromatin binding and enhancer accessibility. Both of these mechanisms appeared to be critical and function as a molecular AND gate for SCC initiation and maintenance, thereby explaining the specificity of the role of ACTL6A amplification in SCCs.

**Highlights:**

- ACTL6A occupancy within BAF complexes is sub-stoichiometric in normal epithelial cells.
- SCC cells upregulate ACTL6A thus increasing ACTL6A assembly with BAF complexes.
- Genome-wide chromatin profiling identifies ACTL6A-dependent regulatory regions.
- Increasing ACTL6A incorporation enhances TEAD-YAP binding to BAF complexes.
- ACTL6A overexpression counteracts polycomb-mediated repression at SCC signature genes.

## Introduction

Mammalian SWI/SNF (also known as BAF) complexes belong to a family of ATP-dependent chromatin remodelers, which contain an ATPase motor that binds nucleosomes and acts to distort or disrupt DNA-histone contacts (Clapier et al., 2017; He et al., 2020; Mashtalir et al., 2020). The enzymatic remodeling activity of the BAF complex is critical for gene regulation allowing specific transcription factors to access their recognition sites (Barisic et al., 2019; Clapier et al., 2017; Kadoch and Crabtree, 2015). Meanwhile, it also facilitates chromatin access for machinery regulating DNA repair, DNA decatenation, and other nuclear processes (Clapier et al., 2017; Kadoch and Crabtree, 2015). BAF complexes are combinatorially assembled from 15 different subunits encoded by 29 genes (He et al., 2020; Kadoch et al., 2013; Mashtalir et al., 2020). They lack sequence-specific DNA recognition; however, subunits of BAF complexes contain domains involved in binding to diverse histone modifications, AT-rich sequences, cruciform DNA structures as well as chromatin and transcriptional regulators that act in concert to guide BAF complex targeting over the genome (Kadoch and Crabtree, 2015). Unique alterations in BAF complex composition during development and in response to signaling further specialize it for engaging specific transcriptional programs (Wu et al., 2009). And yet, the biochemical and structural properties given by individual subunits to diversify the remodeler’s genomic targeting and recruitment of distinct transcriptional regulators remain largely undefined.

The distinct roles of individual subunits in BAF complexes have gained further attention as alterations of different subunits produce specific types of cancers, collectively over 20% of all human cancers (Kadoch et al., 2013; Shain and Pollack, 2013). Frequently, these mutations, including those found on *ARID1A*, are heterozygous and loss-of-function, indicating that BAF subunits are dosage-sensitive and that the complex functions as a tumor suppressor. Dosage-sensitive roles for several of the subunits are also seen in the development of the nervous system and lead to autism and intellectual disability, indicating that the fundamental biochemical mechanisms must also be dosage-sensitive (Ronan et al., 2013; Wenderski et al., 2020). While BAF complex is generally considered a tumor suppressor, some cancers bear gain-of-function BAF alterations, as in synovial sarcomas, where chromosomal translocation at *SS18* results in an oncogenic SS18-SSX fusion, which is present in nearly all cases of synovial sarcoma and appears to be the sole driver mutation (Clark et al., 1994; McBride et al., 2018). This fusion retargets BAF across the genome to reverse polycomb-mediated repression and activate oncogenes including *SOX2* (Kadoch and Crabtree, 2013). Thus, alterations in individual subunits compromise specific biologic actions of BAF complexes, and identifying the underpinning mechanisms holds potential for development of targeted therapy (Kadoch and Crabtree, 2015; Mashtalir et al., 2020; Wilson and Roberts, 2011).

The BAF-subunit gene encoding actin-like 6a (*ACTL6A*, originally called *BAF53A* (Zhao et al., 1998) is located on human chromosomal segment 3q26, an amplification hotspot in multiple SCC types including SCCs in the lung, skin, cervix, and oral mucosa (Ciriello et al., 2013; Heselmeyer et al., 1996; Speicher et al., 1995; Tonon et al., 2005). The amplification develops early in the course of lung SCC development and persists to the metastatic stage, and is thus deemed critical to both tumor initiation and progression (Jamal-Hanjani et al., 2017). A number of driver genes in this amplicon including *PI3KCA, SOX2*, *TP63* have been identified (Bass et al., 2009; Keyes et al., 2011; Simpson et al., 2015; Watanabe et al., 2014), but the role of *ACTL6A* plays in SCC oncogenesis is less known. SCC tumors arise from epithelial tissues, and in epidermis *ACTL6A* expression appears in basal keratinocytes and wanes as they undergo terminal differentiation (Bao et al., 2013). *ACTL6A* over-expression leads to an expanded basal layer of the epithelium, and conversely, loss of *ACTL6A* induces keratinocyte differentiation (Bao et al., 2013). *ACTL6A* also promotes the proliferation of other adult stem cells including hemopoietic and neural stem cells (Krasteva et al., 2012; Lessard et al., 2007). In head-and-neck SCCs, ACTL6A was found to interact with co-amplified TP63 to co-regulate genes promoting proliferation and suppressing differentiation (Saladi et al., 2017). ACTL6A and β-actin form the actin-related protein (ARP) module in BAF complexes and bind the HAS domain of SMARCA4 (BRG1) or SMARCA2 (BRM) ATPases (He et al., 2020; Mashtalir et al., 2020; Szerlong et al., 2008). In yeast, homologs of ACTL6A, Arp7/9, increase the efficiency of ATP utilization by the yeast SWI/SNF complex (Szerlong et al., 2008). Nevertheless, how *ACTL6A* amplification affects SCC chromatin landscape and whether it is attributable to altered BAF complex biochemical properties and interacting surfaces are still unknown.

Here we find that amplification and overexpression of *ACTL6A*, which occurs in ∼25% of SCCs, increases ACTL6A’s normally unsaturated occupancy within BAF complexes. This prompts genome-wide polycomb redistribution, leading to derepression of genes critical for SCC oncogenesis. In addition, increased ACTL6A incorporation directs BAF complex’s interaction with TEAD-YAP transcriptional regulators and prepares the chromatin landscape for TEAD-YAP chromatin binding. Importantly, using structural guided mutagenesis, we found that mutations of two adjacent hydrophobic amino acids within ACTL6A enhanced the binding between BAF-TEAD/YAP complexes and promoted SCC proliferation, thereby defining a potential druggable target for *ACTL6A*-overexpressed cancers. The dosage sensitivity of ACTL6A’s mechanism implies that therapeutic efforts might be directed to reducing ACTL6A function or stoichiometry.

## Results

### BAF complex alterations across multiple SCC types

To comprehensively assess the mutational burden to all BAF subunits in SCCs, we quantified the frequencies of SCC tumors with mutations, copy number variations, and mRNA expression alterations in all 29 BAF-subunit genes using available data sets from three SCC tissue types in the cBioPortal for Cancer Genomics (Cerami et al., 2012; Gao et al., 2013). As chromosome 3q26, *ACTL6A* amplification was prominent as reported (Ciriello et al., 2013; Heselmeyer et al., 1996; Saladi et al., 2017; Speicher et al., 1995; Tonon et al., 2005), in 41% of lung, 18% of head and neck, and 14% of cervical SCCs (24.3% on average); and nearly 50% of SCC tumors had increased *ACTL6A* expression (69% in the lung, 30% head and neck, and 51% cervical SCCs) (Figure 1A). Another BAF-subunit gene*, BRD9*, was also amplified (in total 10% of SCCs) (Figure 1A), but the amplification of *BRD9,* located on chromosome 5q15, infrequently overlapped with *ACTL6A* amplification, suggesting that either might be sufficient (Figure 1B). Surprisingly, the overall point mutation frequencies of BAF-subunit genes were low in all three SCCs despite their prevalence in other cancer types, including in basal cell carcinoma where 26% of cases harbor deleterious mutations in *ARID1A (BAF250A)* (Bonilla et al., 2016; Kadoch et al., 2013; Shain and Pollack, 2013) (Figure 1A).

**Figure 1.**
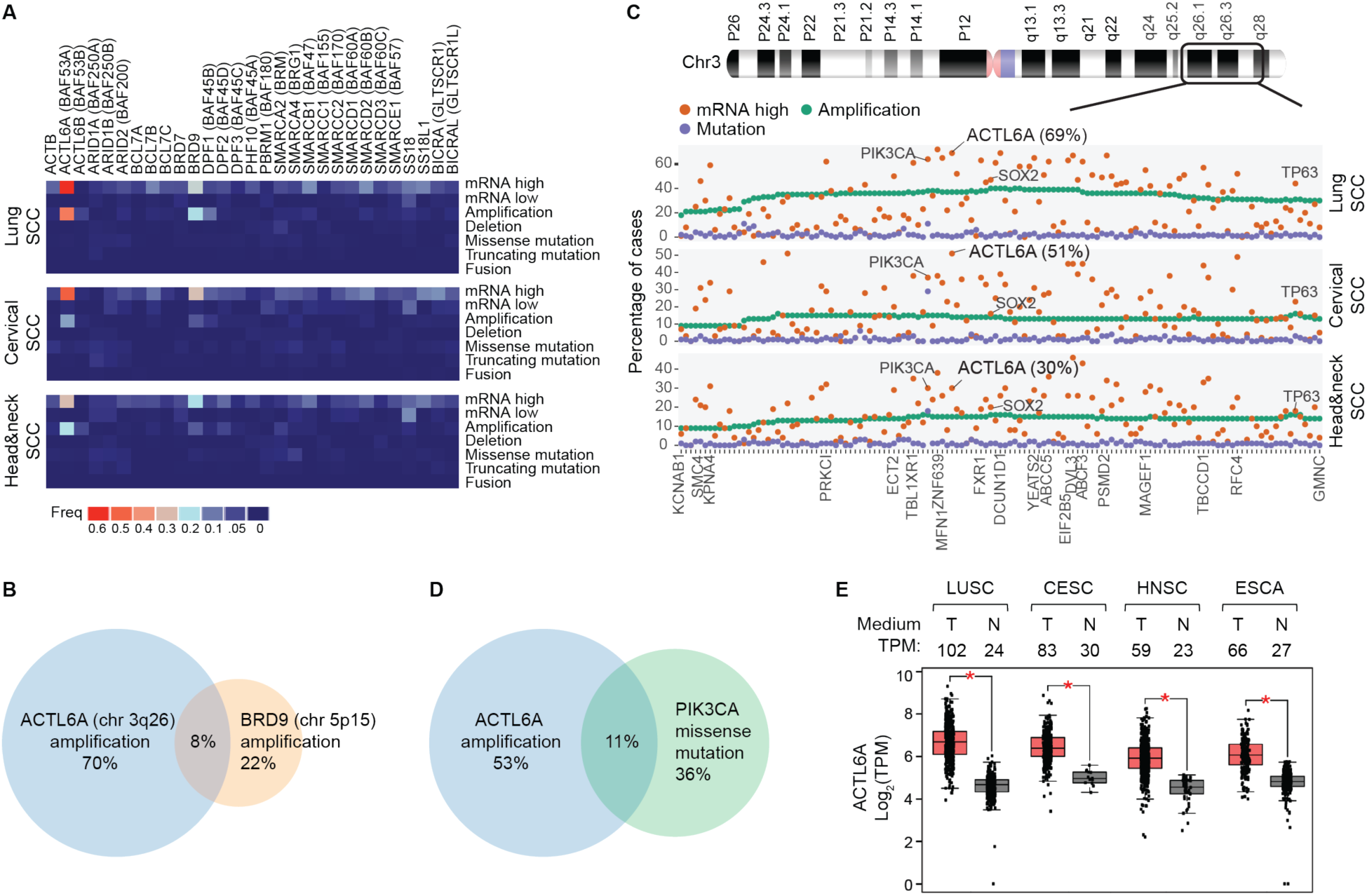
***ACTL6A* amplification and/or overexpression are the most frequent genetic alterations among 29 BAF-subunit genes in lung, head-and-neck and cervical SCCs** (A) Heat maps for alteration frequencies of 29 BAF-subunit genes in three SCC types (cBioPortal). mRNA-high/low: z-score threshold ±2 relative to diploid samples. (B) Venn diagram between SCC tumors with amplification of *ACTL6A* (on chromosome (chr) 3q26), and amplification of *BRD9* (on chr 5p15). Combined cases of lung, cervical and head-and-nect SCCs. (C) Alteration frequencies of 133 genes co-amplified with *ACTL6A* in SCCs. (D) As in (B) for SCC tumors with *ACTL6A* amplification and *PIK3CA* missense mutations. (E) Box plots of *ACTL6A* transcripts-per-million (TPM) in tumors and their paired normal tissues. LUSC: lung SCC. CESC: cervical SCC and endocervical adenocarcinoma. HNSC: head-and-neck SCC. ESCA: esophageal carcinoma. T: tumor samples. N: normal tissue samples. RNA-seq: TCGA and GTEx gene expression data from the GEPIA 2. \**P* < 0.01.

Within the 3q25-28 amplicon, the frequencies of *ACTL6A* upregulation were higher than known SCC drivers *SOX2* and *TP63* and similar to *PIK3CA*, indicating *ACTL6A*’s driver role (Figure 1C). Of note, while mutations in *PIK3CA* and BAF-subunit gene *ARID1A* co-occur to promote ovarian cancer (Chandler et al., 2015), *PIK3CA* mutations in SCCs were frequent but generally exclusive of *ACTL6A* amplification (Figure 1D). The median expression of *ACTL6A* in SCCs was 2- to 4-fold higher than in normal matched tissue samples (4.3-fold in the lung, 2.6-fold in head-and-neck, 2.8-fold in cervical SCCs) (Figure 1E). Overall, contrary to most other cancers, *ACTL6A* amplification and/or over-expression rather than point mutations in BAF complex members is the dominant alteration of BAF complexes in SCCs, implicating a distinct composition and biochemical property of BAF complexes for SCC oncogenesis.

### Increased ACTL6A occupancy with BAF complexes in SCC cells

The dosage-sensitive roles of BAF subunits in neurodevelopment and cancers (Kadoch and Crabtree, 2015) led us to investigate how ACTL6A levels affect BAF complex composition in SCCs. We performed quantitative Western blotting to quantify the number of ACTL6A versus BAF complex molecules per cell in three SCC cell lines bearing over-expressed *ACTL6A*, along with primary normal human keratinocytes (KC; cell type of origin for SCC) (Figures 2A and 2B; Figure S1A). We prepared whole-cell lysates from equal numbers of cells of each cell line, and quantified the amount of a specific protein from a given number of cells using its purified and quantified recombinant proteins for generating a standard curve, which was then used for calculating protein mass and the number of molecules per cell (Figure 2A). The total number of BAF complexes was estimated using an antibody that recognizes both SMARCA4 (BRG1) and SMARCA2 (BRM), which are mutually exclusive catalytic-subunits of BAF complexes. Surprisingly, we found that the number of ACTL6A molecules per normal keratinocyte (111,686±9,850) was only half the number of SMARCA4/SMARCA2 (222,311±21,635 per cell), suggesting that ACTL6A, in fact, is a sub-stoichiometric subunit of the complex in normal keratinocytes (Figure 2B). In all three SCC cell lines we examined, however, ACTL6A molecules were >2-times more numerous than SMARCA4/SMARCA2, which could result in more ACTL6A-containing complexes (ACTL6A: 539,800±33,426 in FaDu (head and neck SCC cell line), 696,016±50,385 in NCI-H520 (lung SCC cell line), 830,683±116,333 in T.T (esophageal SCC cell line); SMARCA4/SMARCA2: 323,542±25,374 in FaDu, 389,563±9,539 in NCI-H520, 315,344±20,536 in T.T) (Figure 2B).

**Figure 2.**
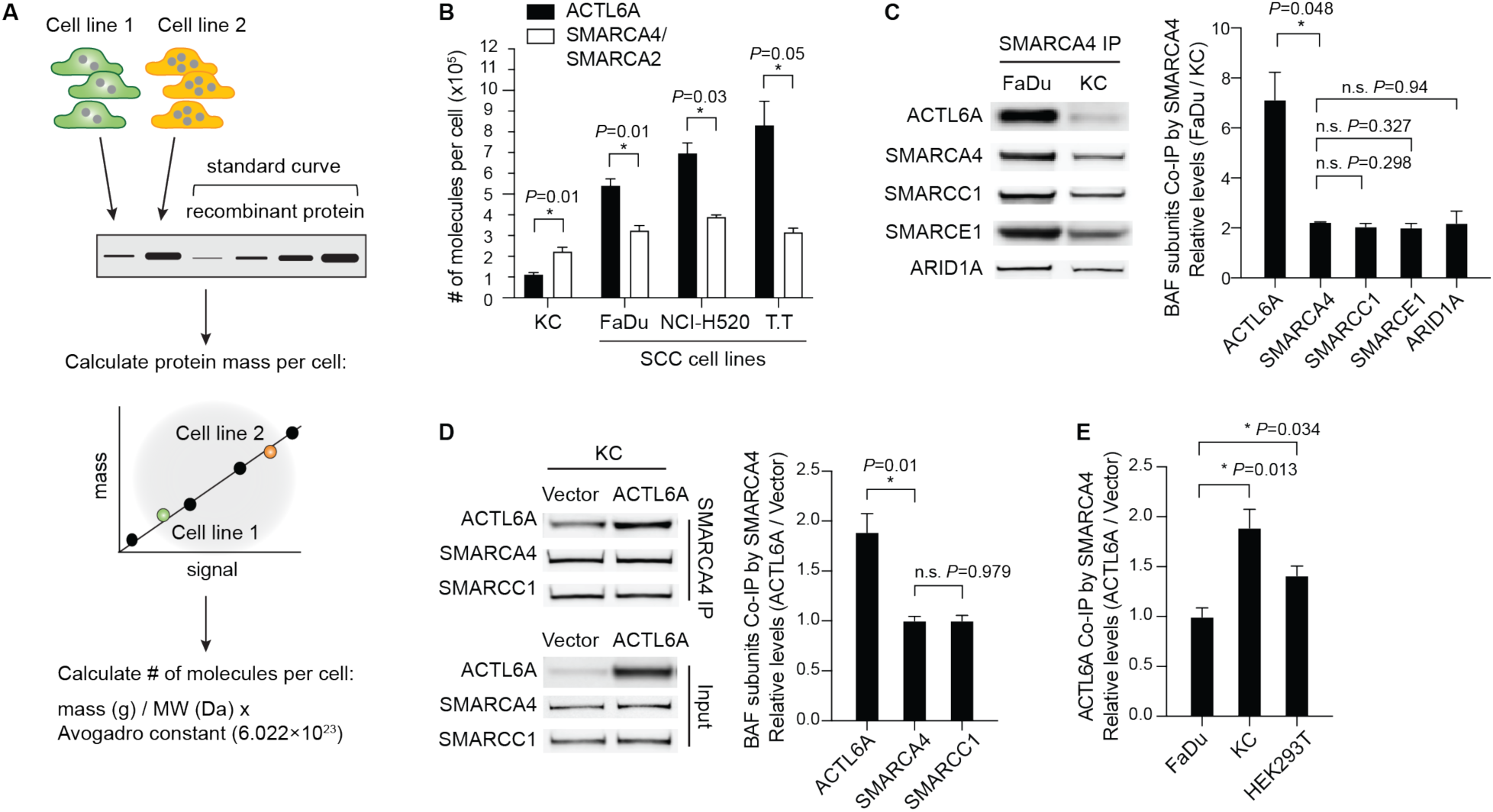
**Increased expression of *ACTL6A* in SCCs drives ACTL6A occupancy within BAF complexes** (A) Outline of method for quantifying the number of molecules of a specific protein per cell from different cell lines. (B) Quantifications of the number of ACTL6A molecules per cell compared to SMARCA4/SMARCA2. SCC cell lines: FaDu (head-and-neck), NCI-H520 (lung), and T.T (esophageal). KC: primary normal human keratinocytes. n=3 experiments. Error bars indicate SEM. \**P* < 0.05. (C) Co-immunoprecipitation (Co-IP) experiments using SMARCA4 antibody. Shown are Western blots and quantifications of relative levels of BAF subunits co-IP’d by SMARCA4 in SCC (FaDu) cells normalized to primary human keratinocytes (KC). n=3 experiments. Error bars indicate SEM. \**P* < 0.05. n.s.: not significant. (D) Co-IP experiments using SMARCA4 antibody in primary human keratinocytes (KC) transduced by lentivirus for *ACTL6A* over-expression and the vector control. Shown are Western blots and quantifications of relative levels of co-IP’d BAF subunits normalized to vector control. n=3 experiments. Error bars indicate SEM. \**P* < 0.05. n.s.: not significant. (E) Quantifications for co-immunoprecipitation (co-IP) experiments by SMARCA4 antibody. Relative levels of co-IP’d ACTL6A in ACTL6A-overexpressing condition normalized to vector-control condition by lentiviral transduction. FaDu: SCC cell line. KC: primary human keratinocytes. HEK293T: human embryonic kidney 293T cells. Error bars indicate SEM. \**P* < 0.05. n=2-3 experiments.

To further compare the occupancy of ACTL6A within BAF complexes in SCC cells to normal keratinocytes, we conducted SMARCA4 immunoprecipitation (Figure 2C). The relative levels (∼7:1) of ACTL6A co-immunoprecipitated by SMARCA4 in SCC cells compared to keratinocytes were substantially higher than of SMARCA4 itself (∼2:1), indicating a specific increase in ACTL6A occupancy within BAF complexes in SCC cells (Figure 2C). In contrast, as for other BAF subunits including SMARCC1 (BAF155), SMARCE1 (BAF57), and ARID1A (BAF250A), their occupancy within the complexes was unaltered in SCC cells, where the relative co-immunoprecipitated levels of those subunits were similar to SMARCA4 (Figure 2C).

To specifically test if ACTL6A can drive complex stoichiometric changes, we reasoned that overexpressing *ACTL6A* in normal cells should increase ACTL6A incorporation into BAF complexes. Indeed, in keratinocytes transduced by lentivirus expressing *ACTL6A*, the levels of ACTL6A co-immunoprecipitated with SMARCA4 antibodies were 1.5- to 2-fold higher than in vector-control cells (Figure 2D). SMARCA4 levels remained unaltered, as did the incorporation of other BAF subunits including SMARCC1 (Figure 2D). Overexpressing *ACTL6A* in another non-SCC cell line HEK293T (human embryonic kidney 293T) also increased ACTL6A occupancy in BAF complexes (Figure 2E). However, elevating ACTL6A levels in SCC cells (FaDu) failed to further its incorporation, indicating the occupancy of ACTL6A within BAF complexes neared saturation in SCC cells (Figure 2E). The increased occupancy was not attributable to the dimerization of ACTL6A as ectopically expressed ACTL6A-V5 did not bind untagged ACTL6A even though both were incorporated into BAF complexes (Figures S1B and S1C). Density sedimentation analysis for SCC cell nuclear extracts showed most ACTL6A proteins co-migrated with SMARCA4 forming a full BAF complex (Figure S1D). Thus, these results reveal an alteration of ACTL6A occupancy within BAF complexes in response to *ACTL6A* amplification or over-expression for SCC development. The occupancy of ACTL6A within the complexes is unsaturated in normal keratinocytes, which becomes saturated by *ACTL6A* overexpression or after amplification in SCCs cells.

### ACTL6A regulates the accessibility of TEAD, POU and FOX transcription factor binding regions over the SCC genome

To identify accessible chromatin regions in the SCC genomes that are dependent on ACTL6A stoichiometry, we conducted ATAC-seq (assay for transposase-accessible chromatin using sequencing) in SCC cells with and without *ACTL6A* knockdown. Reducing ACTL6A levels by siRNA (siACTL6A) did not affect the levels of other BAF subunits including SMARCA4 and SMARCC1, consistent with the notion that ACTL6A is not required for the stability and assembly of BAF complexes (Braun et al., 2021; Krasteva et al., 2012) (Figure S2A). *ACTL6A* knockdown inhibited SCC cell proliferation (Figure S2B), as previously described (Saladi et al., 2017).

*ACTL6A* knockdown in SCC cells caused significant accessibility changes in 4,639 regulatory regions, in which 2,053 displayed decreased and 2,586 displayed increased accessibility (Figure 3A). Interestingly, the top-most significant sequence motifs enriched in regions with decreased accessibility were for TEA domain (TEAD1-4) transcription factors, binding of which appeared to be dependent on the presence of ACTL6A (Figure 3B). The enrichment was further evident as predicted TEAD motifs were found in 818 out of 2053 ACTL6A-dependent accessible regions, whereas these motifs were present in only 219 out of 2586 ACTL6A-repressed regions (Figure 3A). CEBPA, POU-domain and forkhead-box (FOX) motifs were also enriched in ACTL6A-promoted accessible regions, to a lesser extent, whereas CTCF-binding motifs were depleted in these sites, suggesting that insulator elements may be refractory to ACTL6A loss (Figure 3B). 92% of ACTL6A-promoted accessible regions were outside gene promoters by genome annotation analysis, implicating ACTL6A’s role in promoting the accessibility of distal regulatory elements (Figure S2C). In contrast, 20% of ACTL6A-repressed regions were within gene promoters (Figure S2D), which were enriched for transcription factors motifs for AP-1 family members FOS and JUN (Figure S2E), likely reflecting the reaction to genotoxic stress characteristic of BAF subunit depletion (Dykhuizen et al., 2013; Smeyne et al., 1993; Wenderski et al., 2020).

**Figure 3.**
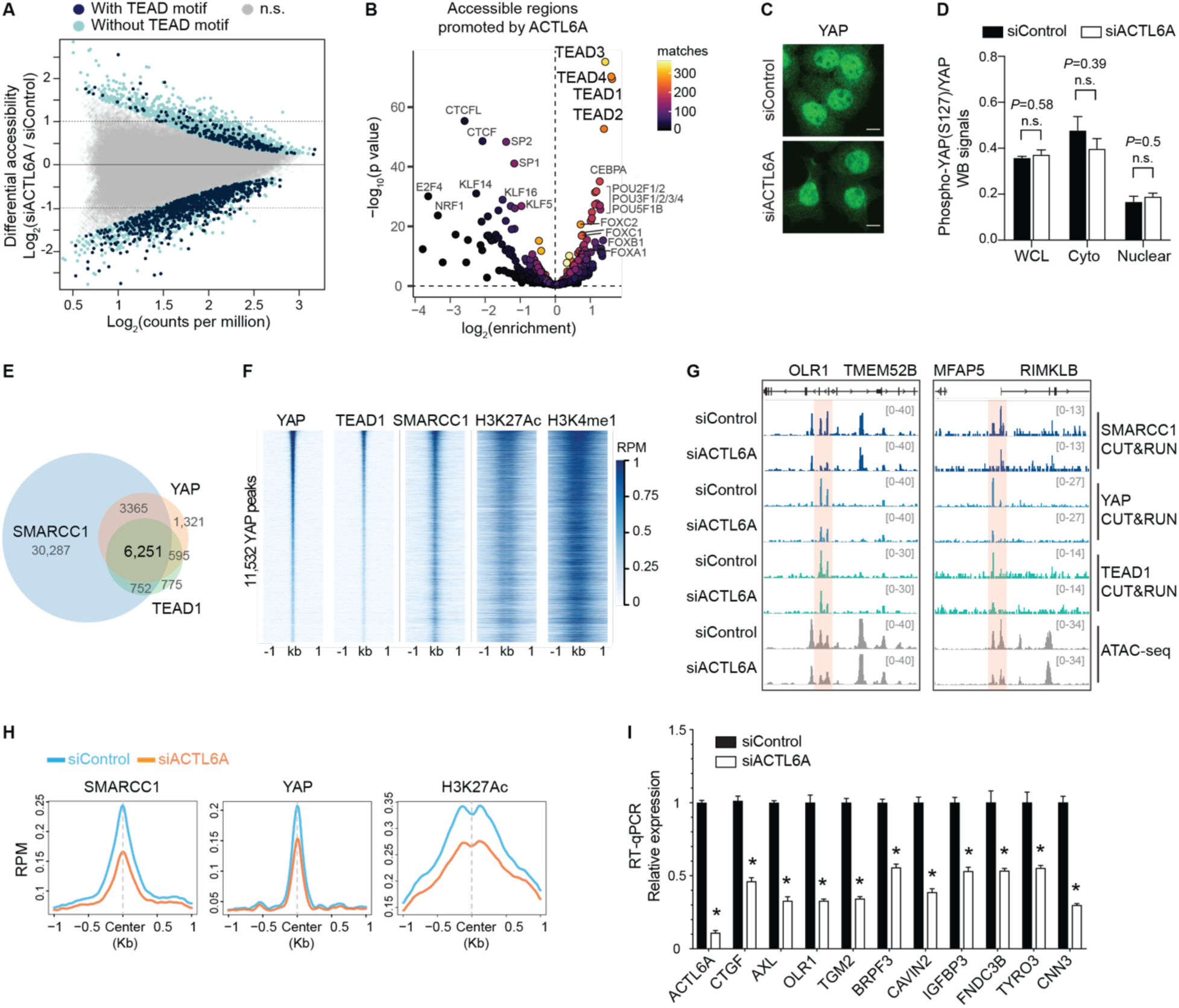
**Genome-wide accessible chromatin and CUT&RUN profiling identifies ACTL6A- dependent regulatory regions in SCC cells** (A) MA plot for ATAC-seq analysis in FaDu SCC cells 72-hours after transfection with *ACTL6A* siRNA (siACTL6A) and control siRNA (siControl). Color coded are significantly altered peaks that contain predicted TEAD-binding motifs (navy) and without TEAD motif (light blue). FDR<0.05. n.s.: not significant. n=2 experiments. (B) Enrichment for predicted transcription-factor (TF) binding motifs in ACTL6A-promoted accessible regions. Matches: number of peaks containing matched TF-binding motifs. (C) Immunofluorescence showing nuclear YAP localization in FaDu SCC cells 72 hours after *ACTL6A* siRNA (siACTL6A) knockdown versus siRNA control (siControl). Scale bars: 10 µm. (D) Quantifications for Western blot (WB) signals of phospho-S127 YAP normalized to total YAP levels. Samples including whole cell lysates (WCL), cytoplasmic extracts (cyto) and nuclear extracts (nuclear) from FaDu cells 72 hours after siRNA transfection. n=3 experiments. (E) Venn diagram showing the overlap of peaks of SMARCC1, YAP and TEAD1 CUT&RUN in FaDu SCC cells. n=2 experiments. (F) Heat maps for CUT&RUN YAP peaks aligned with indicated CUT&RUN peaks. (G) Genome browser tracks showing regions with reduced accessibility (ATAC-seq) and reduced binding of SMARCC1, YAP and TEAD1 (CUT&RUN) upon *ACTL6A* knockdown (siACTL6A) compared to siControl. (H) Profiles of SMARCC1, YAP and H3K27Ac CUT&RUN over ACTL6A-promoted accessible sites. Error bars indicate SEM. \**P* < 0.05. n.s.: not significant. (I) RT-qPCR showing YAP/TEAD target genes regulated by ACTL6A. n=3 experiments. Error bars indicate SEM. \**P* < 0.05.

TEADs form complexes with transcriptional co-activators YAP/TAZ and they act downstream of multiple signaling pathways including mechanotransduction and the Hippo pathway critical for tumorigenesis, organ size control, and regeneration (Yu et al., 2015a; Zanconato et al., 2016). In mammalian skin, YAP/TAZ promotes SCC initiation and progression (Debaugnies et al., 2018; Schlegelmilch et al., 2011; Vincent-Mistiaen et al., 2018). In *Drosophila*, Hippo signaling has been shown dependent on BRM and the fly SWI/SNF (or called BAP) complex (Jin et al., 2013; Oh et al., 2013). However, the underlying mechanism remains elusive. Previous studies suggest ACTL6A loss inhibits YAP activity by upregulating the WWC1 gene, which encodes a scaffold protein in the Hippo pathway that promotes YAP retention in the cytoplasm (Saladi et al., 2017). However, we did not observe changes in YAP subcellular localization by immunostaining in siACTL6A SCC cells, where most YAP proteins remained in the nucleus as in siControl cells (Figure 3C). Western blotting also did not show changes in the phosphorylation of YAP S127, which promotes its cytoplasmic retention (Yu et al., 2015a) (Figure 3D), and WWC1 expression remained unaltered upon ACTL6A knockdown (Figure S2F). Thus, changes seen in the accessibility of TEAD enhancers upon *ACTL6A* loss were not due to YAP nuclear translocation in these cell lines. Instead, it implied that ACTL6A-BAF complexes directly promote the remodeling of local chromatin at TEAD enhancers.

If the accessibility of TEAD enhancers were directly modulated by ACTL6A-BAF complexes, ACTL6A-BAF complexes should co-bind with TEAD-YAP across the genome. To test this prediction, we conducted CUT&RUN to profile the distribution of TEAD1, YAP, and BAF complexes (SMARCC1, a DNA-binding subunit of BAF complexes) genome-wide. As expected, the regions bound by TEAD1 and YAP largely overlapped (Figures 3E and 3F). Remarkably, 91% of TEAD1-YAP co-bound regions were also bound by SMARCC1, indicating their co-localization on chromatin (Figures 3E and 3F). Furthermore, 79% of YAP/TEAD1/SMARCC1 co-bound regions were at active enhancers marked by the histone modifications H3K27Ac and H3K4me1, concordant with earlier observations of TEAD-YAP binding at active enhancers (Stein et al., 2015; Zanconato et al., 2015) (Figure 3F; Figure S3A). A *de novo* motif search identified TEAD motifs in the 6,251 shared peaks, confirming the specificity of YAP and TEAD1 CUT&RUN profiling (Figure S3B). Notably, while BAF complex-bound regions were enriched most for binding motifs of FOSL2 (JUNB) and SP2 (Figure S3C), the accessibility decreases upon ACTL6A loss were largely at TEAD motifs (Figures 3A and 3B), indicating a specific effect of ACTL6A in the complex on promoting TEAD chromatin binding. CUT&RUN profiling further confirmed the presence of YAP and TEAD1 at ACTL6A-promoted accessible regions marked by enhancer mark H3K4me1, in contrast to low YAP and TEAD1 levels at ACTL6A- repressed regions, which spanned H3K4me3-marked promoters (Figures S3D, S2C and S2D). *ACTL6A* knockdown reduced the binding of SMARCC1 and YAP-TEAD1 at enhancers that also lost accessibility, accompanied by reduced levels of the active mark H3K27Ac (Figures 3G and 3H). Thus, these results indicate that BAF complexes and TEAD-YAP co-bind across the genome, and ACTL6A functioning within the BAF complex plays a direct and specific role in targeting BAF complexes to TEAD-YAP enhancers and preparing the chromatin landscape to allow TEAD-YAP chromatin binding and transcription activation at their target loci.

The decreases in the accessibility at predicted TEAD motifs were accompanied by reduced expression of TEAD/YAP/TAZ target genes in *ACTL6A*-knockdown SCC cells (Figure 3I; Figure S3E). Using RNA-seq analysis, we identified 188 differentially expressed genes between siACTL6A and siControl conditions in at least two of the three SCC cell lines, which included previously identified TEAD/YAP/TAZ target genes (Zhang et al., 2009) (Figures S3E and S3F). These targets were validated by quantitative PCR (qPCR) of independently prepared samples (Figure 3I). Notably, target genes of *TP63* and *SOX2* (Watanabe et al., 2014), known SCC drivers that are co-amplified with *ACTL6A* (Figure 1B), were largely unaltered (Figure S3G), suggesting that *ACTL6A* is not essential for the downstream transcriptional programs of *TP63* and *SOX2* despite their co-amplification with *ACTL6A* in SCCs. It is possible that the pioneer factor property of SOX2, which can initiate chromatin opening (Dodonova et al., 2020), might account for its refractory to ACTL6A, or the remodeling activity from residual BAF complexes is sufficient for SOX2’s chromatin binding.

### Increasing ACTL6A levels induces TEAD-YAP binding to BAF complexes

The dependency of TEAD cognate motif accessibility and chromatin binding upon ACTL6A levels suggested that an interaction of TEAD and its co-activator YAP with BAF complexes and ACTL6A incorporation might modulate the interaction. To test whether ACTL6A assembled within the BAF complex serve to directly recruit TEAD-YAP, we conducted immunoprecipitation experiments. Using TEAD4 and pan-TEAD antibodies, we were able to co-immunoprecipitate SMARCA4 and several other BAF subunits from nuclear extracts of SCC cells (Figure 4A; Figure S4A). Furthermore, the reciprocal immunoprecipitation with SMARCA4 and ACTL6A antibodies yielded TEAD proteins (Figures S4B and S4C). Antibodies to YAP also co-immunoprecipitated BAF subunits (Figure 4B). Thus, TEAD-YAP directly interact with BAF complexes.

**Figure 4.**
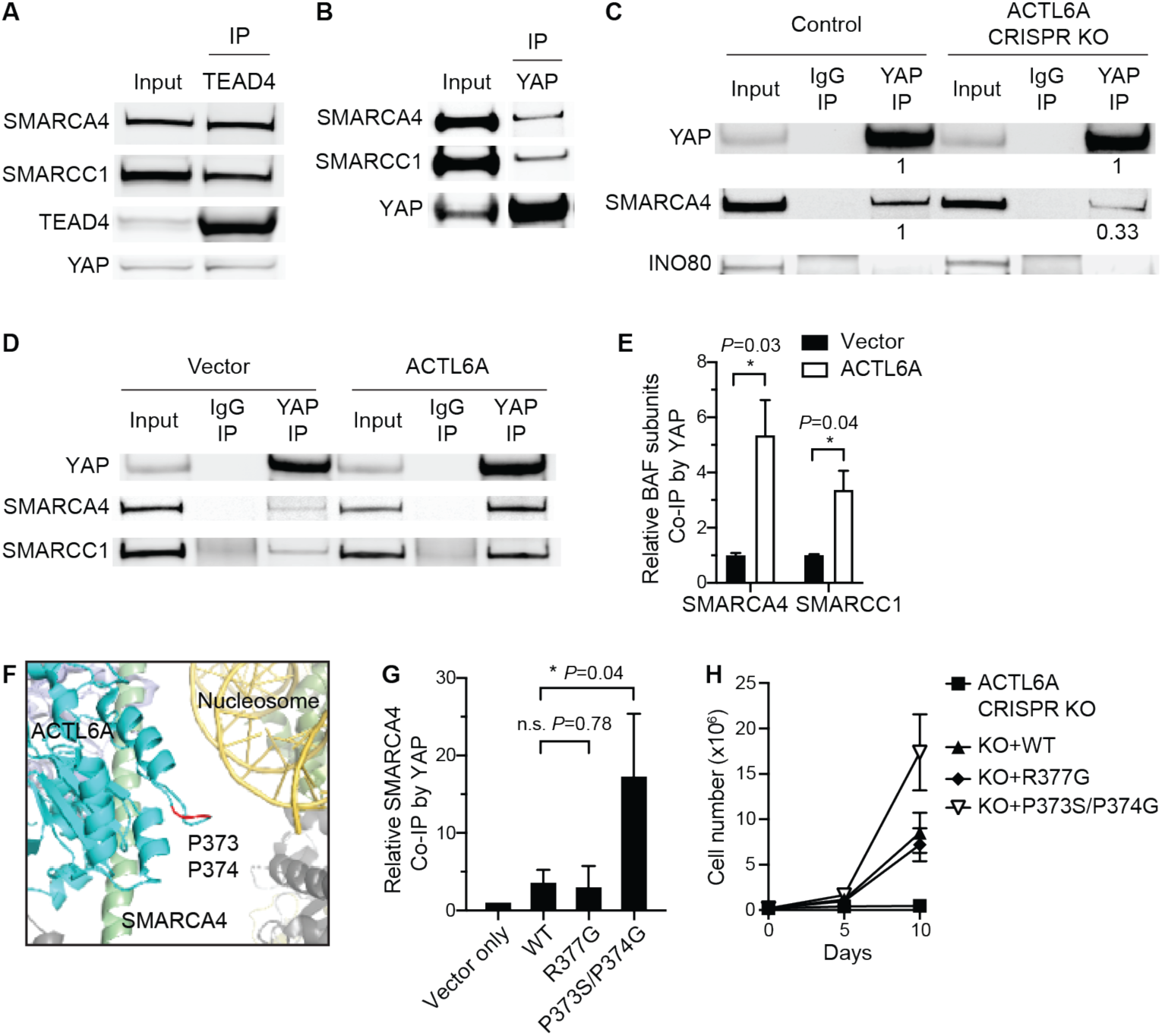
**ACTL6A promotes the direct binding of TEAD-YAP to BAF complexes** (A-B) Co-IP experiments by TEAD4 (A) and YAP (B) antibodies using nuclear extracts from FaDu SCC cells. (C) Co-IP experiments by YAP and IgG antibodies showing decreased binding of SMARCA4 with YAP in *ACTL6A* CRISPR-knockout (KO) FaDu SCC cells. Quantifications normalized to control. INO80: INO80- complex subunit. (D) Co-IP experiments by YAP and IgG antibodies in primary human keratinocytes overexpressing *ACTL6A* and vector-control. (E) Quantifications for (D) showing increased binding of YAP and BAF subunits upon *ACTL6A* over-expression. Normalized to vector control. n=3 experiments. Mean ± SEM. \**P* < 0.05. (F) The human BAF complex cryo-EM structure (PDB: 6LTJ) showing the position of ACTL6A P373/P374 residues (marked in red) in the nucleosome-bound BAF complex. Cyan: ACTL6A. Red: P373/P374 residues of ACTL6A. Green: HSA domain of SMARCA4. Yellow: Nucleosomal DNA. Olive: histone octamer. (G) Quantifications for co-IP experiments by YAP antibodies in keratinocytes overexpressing WT, R377G and P373S/P374G ACTL6A. Normalized to vector control. n=3 experiments. Mean ± SEM. \**P* < 0.05. n.s.: not significant. (H) Growth curves of FaDu SCC cells transduced with lentiviral constructs for *ACTL6A* CRISPR-KO and simultaneously reconstituted with KO-resistant WT, R377G, P373S/P374G ACTL6A, or vector-control. n=3 experiments. Error bars indicate SD.

To examine whether the interaction of TEAD-YAP and BAF complexes is dependent on ACTL6A, we conducted ACTL6A loss-of-function and gain-of-function analyses. Reducing ACTL6A levels in SCC cells by siRNA or CRISPR/Cas9 blocked the binding of BAF complexes to YAP (Figure 4C; Figure S4D). Remarkably, overexpressing *ACTL6A* in normal human keratinocytes, which enhanced ACTL6A occupancy within BAF complexes (Figure 2D), caused increased binding of BAF complexes with YAP and TEADs (Figures 4D and 4E; Figure S4E). We did not observe YAP binding to INO80, another ACTL6A- associated chromatin remodeler (Figure 4C). Thus, ACTL6A is necessary and sufficient for the interaction between BAF complexes and TEAD-YAP in SCC cells.

The WW domains of YAP recognize the PPxY motif on its interaction partners (Chen and Sudol, 1995). While ACTL6A does not contain a PPxY motif, we speculated a proline-rich loop structure (PPSMRLKLI; a.a. 373-381) on ACTL6A that extends toward the nucleosomal DNA bound by the BAF complex (He et al., 2020) may serve as an alternative interaction mechanism (Figure 4F). To examine the effect of ACTL6A P373S/P374G double mutations on the interaction between YAP and BAF complexes, we introduced P373S/P374G ACTL6A in human keratinocytes with WT ACTL6A as control and conducted YAP immunoprecipitation. While WT ACTL6A overexpression is sufficient to enhance the interaction (Figures 4D and 4E), unexpectedly, we found that overexpression of P373S/P374G ACTL6A induced higher levels of YAP binding to BAF complexes than WT ACTL6A (Figure 4G). A nearby R377G mutation did not show this effect on BAF-YAP binding (Figure 4G). To examine whether the increased binding might promote SCC proliferation, we knocked out endogenous *ACTL6A* using CRISPR/Cas9 and simultaneously reconstituted with WT or mutated ACTL6A. Reconstituting WT ACTL6A rescued the proliferation defects caused by *ACTL6A* knockout, and notably, P373S/P374G mutants promoted SCC growth better than WT or R377G ACTL6A (Figure 4H). These mutations neither compromised ACTL6A stability nor altered its incorporation into the BAF complex (Figure S4F). Together, these results indicate that ACTL6A regulates TEAD-YAP activity to drive SCC growth by producing interaction surfaces on BAF complexes for TEAD-YAP binding.

### ACTL6A over-expression redistributes polycomb over the genome

The SWI/SNF and polycomb-repressive complexes (PRCs) play opposing roles in epigenetic regulation. In flies, loss-of-function mutations in the SWI/SNF subunits suppress defects that are conferred by mutations in PRC1 and rescue the developmental phenotypes of PRC1 mutants, indicating a rather dedicated relationship between these two classes of chromatin regulators (Tamkun et al., 1992). In synovial sarcoma and several other BAF-subunit mutated cancers, PRCs are important primary targets of mammalian SWI/SNF or BAF complexes and distinct mutations in BAF complexes can result in either a gain or loss of its ability to evict PRCs (Ho et al., 2011; Kadoch and Crabtree, 2013; Kadoch et al., 2017; Kia et al., 2008; Stanton et al., 2017). To determine early consequences of *ACTL6A* amplification and whether they are attributable to perturbation of the BAF-PRC balance, we examined by CUT&RUN the genome-wide distribution of PRC2-modified histone mark H3K27me3 in primary normal human keratinocytes following *ACTL6A* over-expression.

Remarkably, *ACTL6A* over-expression led to H3K27me3 redistribution over the genome (Figure 5A). We identified gene promoters with altered H3K37me3 deposition, and most of them showed decreased H3K27me3 levels, consistent with BAF’s ability to rapidly and directly evict PRC by an ATP-dependent mechanism (Kadoch et al., 2017; Stanton et al., 2017) (Figure 5B). Affected promoters were primarily bivalent, i.e. marked by both H3K27me3 and H3K4me3, supporting BAF’s role in maintaining bivalent chromatin state (Stanton et al., 2017) (Figures 5B and 5C). To explore how changes in H3K27me3 levels affected transcription, we conducted RNA-seq analysis. As expected, alterations in H3K27me3 and transcription were largely negatively correlated. Genes with reduced H3K27me3 tended to have increased transcription, while genes with increased H3K27me3 tended to display decreased expression (Figure 5C). Notably, a subset of genes with altered H3K27me3 levels lacked significant changes in their transcription, suggesting that H3K27me3 alterations induced by *ACTL6A* over-expression were likely direct consequences rather than secondary effects from transcriptional changes.

**Figure 5.**
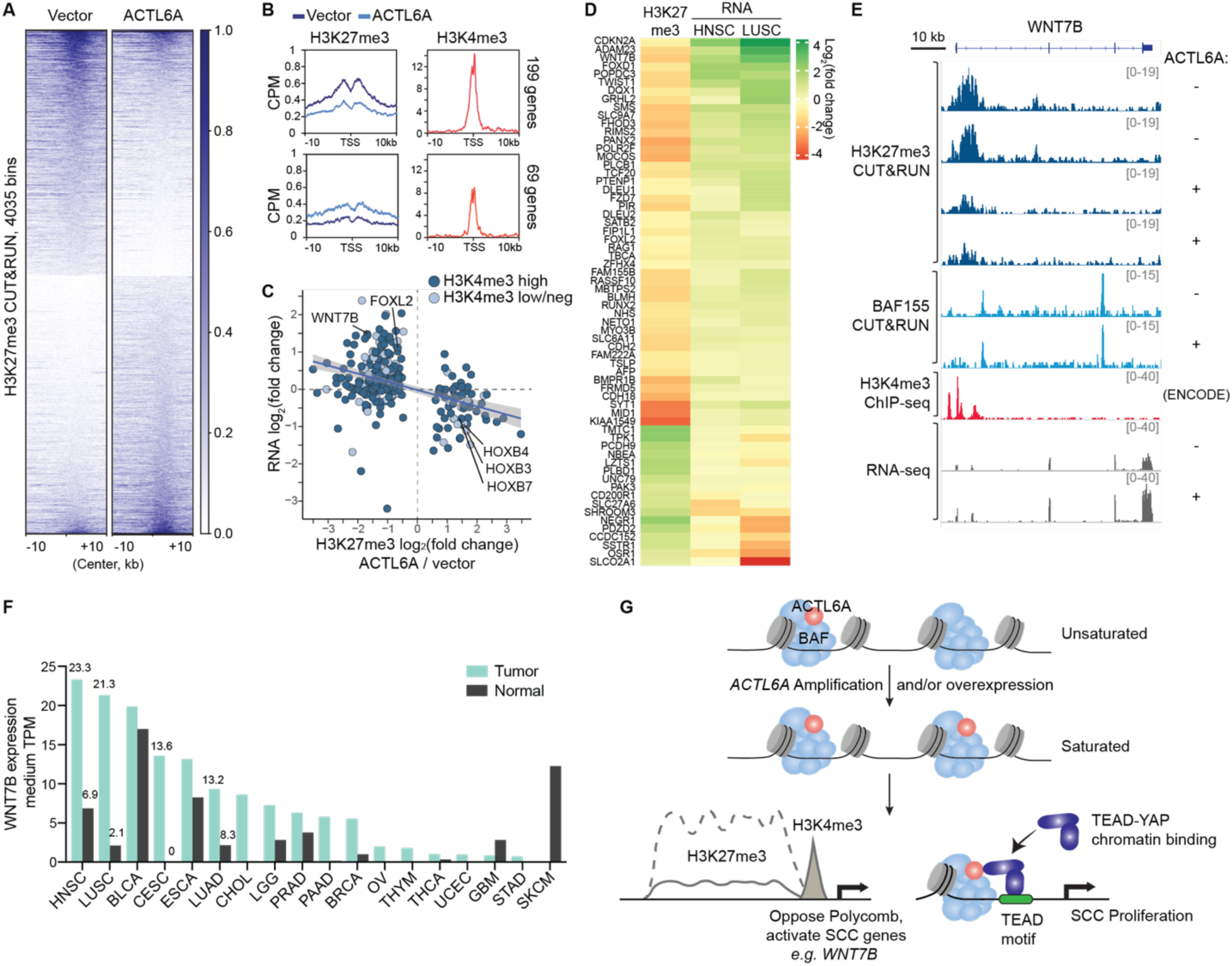
***ACTL6A* over-expression leads to redistribution of H3K27me3 and activation of SCC genes** (A) Heat maps for H3K27me3 CUT&RUN differential 5kb-bins between primary human keratinocytes overexpressing *ACTL6A* and vector-control. Scale: counts per million, CPM. n=2 experiments. (B) Profiles over TSS with decreased (top) and increased (bottom) H3K27me3 levels upon *ACTL6A*- overexpression versus vector control by CUT&RUN. Right: H3K4me3 ChIP-seq profiles of human keratinocytes (ENCODE). CPM: counts per million. (C) Scatterplot of H3K27me3 and RNA fold-changes for genes with differential H3K27me3 levels upon *ACTL6A*-overexpression. Color codes: H3K4me3 levels, high versus low or negative (neg). (D) Heatmap showing ACTL6A-dependent PRC target genes with corresponding transcriptional changes in SCC tumors. H3K27me3: CUT&RUN as in (A). RNA: HNSC (head-and-neck SCC) and LUSC (lung SCC) tumor versus normal tissue from GEPIA 2. (E) Genome browser tracks at bivalent *WNT7B* gene upon *ACTL6A*-overexpression (+) compared to vector control (-). (F) *WNT7B* medium expression levels in tumors and paired normal tissues across various cancers. Data from GEPIA 2. TPM: transcripts per million. HNSC: head-and-neck SCC. LUSC: lung SCC. BLCA: bladder urothelial carcinoma. CESC: cervical SCC and endocervical adenocarcinoma. ESCA: esophageal carcinoma. LUAD: lung adenocarcinoma. CHOL: cholangiocarcinoma. LGG: brain lower grade glioma. PRAD: prostate adenocarcinoma. PAAD: pancreatic adenocarcinoma. BRCA: breast invasive carcinoma. OV: ovarian serous cystadenocarcinoma. THYM: thymoma. THCA: thyroid carcinoma. UCEC: uterine corpus endometrial carcinoma. GBM: glioblastoma multiforme. STAD: stomach adenocarcinoma. SKCM: skin cutaneous melanoma. (G) Model for *ACTL6A*-amplification driven oncogenic mechanism in SCCs. Amplification or overexpression of *ACTL6A* leads to full occupancy of BAF complexes giving rise to two mechanisms promoting SCC initiation and maintenance.

The decrease of H3K27me3 marks upon *ACTL6A* over-expression was not due to keratinocyte differentiation, which in contrast leads to a global loss of H3K27me3 and an accompanying decrease in *ACTL6A* (Bao et al., 2013; Ezhkova et al., 2009). Furthermore, the H3K27me3 levels at polycomb-repressed differentiation genes such as *KRT1* and *LOR* were unaltered (Figure S5A). Also unchanged were the expression of keratinocyte differentiation genes (*KRT1, KRT10, IVL, LOR*) and progenitor markers (*KRT14, KRT5, TP63*) (Figure S5B). Consistent with the observation that conditional loss of BAF subunits decreases H3K27me3 levels in *HOX* clusters (Ho et al., 2011), we observed a gain of H3K27me3 in the *HOXB* locus upon *ACTL6A* over-expression (Figure 5C; Figure S5C). Thus, over-expression of *ACTL6A* in normal human keratinocytes leads to a redistribution of H3K27me3 over the genome.

Because ACTL6A amplification is a very early event in the pathogenesis of SCC (Jamal-Hanjani et al., 2017), we reasoned that overexpressing it in normal keratinocytes might initiate a program of SCC gene expression. If PRC ejection were a major driving mechanism, then these SCC genes should be distinguished by PRC loss upon ACTL6A expression. To determine whether the polycomb target genes affected by ACTL6A dosage are also misregulated in SCC tumors *in vivo*, we examined their transcripts in SCC tumors versus normal tissues using CGA/GTEx data sets available in GEPIA (Tang et al., 2017) (Figure 5D). Interestingly, we found 64 of the PRC targets displayed corresponding changes in the RNA levels in either lung SCC (LUSC) or head-and-neck SCC (HNSC) tumors compared to paired normal tissues with *p*-value<0.05. In total 47 genes were preferentially upregulated in LUSC or HNSC tumors that had reduced H3K27me3 levels upon *ACTL6A* over-expression, whereas only 17 genes had decreased expression in LUSC or HNSC tumors with increased H3K27me3 levels by *ACTL6A* gain (Figure 5 D). The derepressed genes included *WNT7B* (Figures 5E and 5F). *WNT7B* encodes a Wnt ligand and has been found to contribute to skin carcinogenesis in *Cdkn2ab* knockout mouse model (Krimpenfort et al., 2019) and promote proliferation and invasion of oral SCC cells (Shiah et al., 2014). In pancreatic adenocarcinoma, *WNT7B* promotes tumor’s anchorage-independent growth and sphere formation (Arensman et al., 2014). *ACTL6A* overexpression induced *WNT7B* upregulation accompanied with reduced H3K27me3 at its bivalent promoter marked with H3K4me3 (Figure 5E). Two BAF155 CUT&RUN peaks near the H3K27me3 domain and in *WNT7B* gene body were unaltered, suggesting that ACTL6A incorporation into BAF complex did not affect BAF chromatin binding but instead affects its activity in antagonizing PRCs (Figure 5E). Upregulation of *WNT7B* occurred in several types of SCCs including head-and-neck SCC (HNSC) and lung SCC (LUSC), as well as cervical squamous cell carcinoma and endocervical adenocarcinoma (CESC) and esophageal carcinoma (ESCA) (Figure 5F).

Besides *WNT7B*, other ACTL6A-dependent PRC targets known to play roles in SCC oncogenesis included *TWIST1*, which is associated with epithelial–mesenchymal transition (EMT) in esophageal and head-and-neck SCCs (Jouppila-Matto et al., 2011; Lee et al., 2012); and *SATB2* (special AT-rich binding protein 2), which promotes survival and chemoresistance of head-and-neck SCC cells in part by interacting with **Δ**Np63*α* (Chung et al., 2010) and also drives carcinogenesis of oral SCC, colorectal cancer and osteosarcoma (Ge et al., 2020b; Seong et al., 2015; Yu et al., 2017) (Figure 5D). Several other ACTL6A- dependent PRC target genes belong to the forkhead box (FOX) family, members of which are often repressed by polycomb and poised for activation (Golson and Kaestner, 2016) (Figure 5D). *FOXD1* upregulation induces EMT and chemoresistance of oral SCC cells (Chen et al., 2020); and *FOXL2* has been found to be upregulated in SCC tumors (Ge et al., 2020a), and is a driver of granulosa-cell tumor (Shah et al., 2009). Another ACTL6A-dependent PRC target, *CDKN2A* (Figure 5D), is considered a tumor suppressor; however, over-expression of *CDKN2A* has been noted in several tumors including SCC tumors (Romagosa et al., 2011). In sum, our results suggest that before malignant transformation, early *ACTL6A* over-expression in epithelial cells is sufficient to reduce polycomb-mediated repression of genes for SCC oncogenesis by perturbing chromatin architecture and BAF-PRC opposition.

## Discussion

Our studies reveal that *ACTL6A* gene amplification and/or over-expression leads to its increased occupancy within BAF complexes that then facilitates the establishment of an altered chromatin state for SCC development (Figure 5G). BAF subunits are highly dosage-sensitive (Kadoch and Crabtree, 2015). Mutations of BAF subunit genes implicated in human cancers and neurological disorders such as autism and intellectual disability are commonly heterozygous, indicating that a half-normal level is biologically significant (Kadoch and Crabtree, 2015). Thus, the 2- to 5-fold increase in *ACTL6A* levels in SCCs and the consequent increase in ACTL6A occupancy within the complex and proliferation are consistent with the dosage-sensitive role for BAF complexes in human diseases. Our studies indicate that the increased dosage of ACTL6A has two consequences. First, complexes having stoichiometric occupancy of BAF are more effective at evicting polycomb over the genome; and secondly, BAF’s function prepares the genome to receive Hippo signals mediated by TEADs and YAP (Figure 5G). Our studies indicate that both mechanisms are critical and likely to function as an epigenetic AND gate for SCC initiation and maintenance. The requirement of both ACTL6A-dependent mechanisms likely explains the oncogenic specificity of ACTL6A amplification.

In the development of the mammalian nervous system, ACTL6A exchanges with ACTL6B to generate neuron-specific nBAF complexes that coordinate gene expression underlying cell cycle exit and the initiation of neural differentiation (Braun et al., 2021; Lessard et al., 2007). In epithelial cells, ACTL6A levels fall as the cells differentiate. This reduction triggers a switch from proliferation to keratinocyte differentiation with the activation of keratinocyte differentiation genes including *KLF4* (Bao et al., 2013; Krasteva et al., 2012; Lu et al., 2015). ACTL6A is also essential for proliferation and maintaining stem cell potency (Bao et al., 2013; Krasteva et al., 2012). Thus, we propose that the degree of occupancy of ACTL6A in chromatin remodelers is regulatory and therefore its dosage acts as a decisive signal underlying the transition between chromatin states during the initiation of SCCs.

Intriguingly, in contrast to *ACTL6A* amplification in SCCs, basal cell carcinoma (BCC), another cancer originating from basal epithelial cells, has a high frequency of heterozygous loss-of-function mutations in the BAF-subunit *ARID1A* (Bonilla et al., 2016), while such mutations are rare in SCCs (Figure 1A). This indicates different epidermal lineages (Sanchez-Danes and Blanpain, 2018) are specifically susceptible to distinct BAF complex alterations and illustrates the biologic specificity of their functions. Recent structural studies (He et al., 2020; Mashtalir et al., 2020) suggest that this specificity emerges from combinatorial assembly of the products of 29 genes encoding the 15 subunits creating composite surfaces at their interfaces available to interact with proteins such as TEADs and YAP.

Considerable effort has been dedicated to developing TEAD/YAP/TAZ small molecule inhibitors, with limited success (Calses et al., 2019). We find that ACTL6A incorporation promotes TEAD-YAP binding to BAF complexes, which leads to enhanced TEAD-YAP chromatin binding and transcriptional activity, suggesting a new therapeutic approach. The activating effects of ACTL6A as a BAF subunit on the TEAD-YAP pathway are consistent with genetic studies in flies, in which mutations in Brahma, the fly homolog of the SWI/SNF ATPase, hinder transcriptional activation by Yorkie (YAP homolog) (Jin et al., 2013; Oh et al., 2013). Our findings also support studies in breast epithelial lineage commitment where BAF complexes interact with the Hippo pathway component TAZ and positively regulate TAZ-induced transcription (Skibinski et al., 2014). Our CUT&RUN genome-wide mapping of BAF complex and TEAD-YAP elucidate their interaction across the genome and reveal the dependency of TEAD-YAP enhancer accessibility on ACTL6A-containing BAF complexes, which hence prepare the chromatin landscape to allow TEAD-YAP mediated transcription at their target loci. Of note, others (Saladi et al., 2017) reported ACTL6A activates YAP rather by an indirect mechanism, wherein ACTL6A, in collaboration with TP63, controls YAP nuclear localization by repressing genes including WWC1 that modulate YAP nuclear-cytoplasm shuttling. However, we do not detect changes in YAP subcellular localization upon ACTL6A loss; instead, we find BAF complexes directly interact with YAP and TEAD, and the interaction is dependent on ACTL6A.

The failure of YAP pathway inhibition to provide therapeutic benefit (Calses et al., 2019) may be related to our finding that ACTL6A uses another mechanism to drive SCC oncogenesis. We discovered that *ACTL6A* over-expression resulting from either amplification or other means is sufficient to induce polycomb eviction and redistribution, resulting in the activation of genes known to have roles in SCCs. Although the exact mechanism of BAF-PRC antagonism remains unknown, the effects could be rooted in altered SMARCA4/SMARCA2 ATPase activity, which have been shown to be required for BAF complexes to evict PRCs (Kadoch et al., 2017; Stanton et al., 2017) and in yeast, is promoted by the ACTL6A homologs Arp7/9 (Szerlong et al., 2008). Interestingly, the outcomes of ACTL6A-induced polycomb redistribution are rather selective for bivalent genes such as *WNT7B*, the roles of which in tumor initiation merit further investigation given the early occurrence of *ACTL6A* amplification during SCC development (Jamal-Hanjani et al., 2017), as does the roles of ACTL6A in the interplay between BAF and PRC complexes. Of note, despite its high mutation rate, *CDKN2A* is over-expressed in some SCC tumors (Romagosa et al., 2011) and the reduction of polycomb repression at *CDKN2A* upon *ACTL6A* over-expression might reflect the specific pathways of oncogenesis in SCCs. In line with this, polycomb removal and *CDKN2A* activation have also been shown as a direct outcome of gain of BAF complex activity upon *SMARCB1* (or called *SNF5*) re-expression in *SMARCB1*-deficient tumor cells (Kia et al., 2008).

In summary, our studies demonstrate that stoichiometry within a chromatin regulatory complex can be oncogenic, and that the dynamics of ACTL6A occupancy in these complexes may play roles in normal development by enabling protein-protein interactions with key regulators engaging in proliferation and stem cell function. Our studies indicate that both polycomb redistribution and TEAD-YAP facilitation are essential downstream mechanisms for the initiation and maintenance of SCCs. Therefore, therapeutic efforts might be directed to reducing ACTL6A function or stoichiometry. The discovery that mutations of two adjacent residues in ACTL6A enhance proliferation, provide a precise therapeutic target.

## Acknowledgments

We thank all Crabtree laboratory members for intellectual input and suggestions; B. Keyes, S. Lin and W. Lu for critical reading of the manuscript; S. HeniKoff for providing protein A-MNase (pA-MN) for CUT&RUN experiments; X. Xiong and T. Chai for providing reagents. C.-Y.C. was supported by the GSK Sir James Black fellowship. Z.S. is supported by EMBO Long-Term Fellowship EMBO ALTF 1119-2016 and by Human Frontier Science Program Long-Term Fellowship HFSP LT 000835/2017-L. G.R.C. is an investigator of the Howard Hughes Medical Institute. This work was supported by the Howard Hughes Medical Institute, NIH grant CA163915, Department of Defense grant W81XWH-16-1-0083 to G.R.C and the David Korn Professorship. K.M.L. is a Packard Foundation Fellow, Pew Scholar, Human Frontiers Science Program Young Investigator and the Anthony DiGenova Endowed Faculty Scholar; and was supported by the Stanford Ludwig Center for Cancer Stem Cell Research. W.J.G is a Chan Zuckerberg Biohub investigator and acknowledges grant nos. 2017-174468 and 2018-182817 from the Chan Zuckerberg Initiative.

## Author Contributions

G.R.C. and C.-Y.C. conceived the project, designed the experiments and wrote the manuscript. C.-Y.C. conducted all experiments. Z.S. performed the bioinformatic analyses and assisted experiments. A.K. assisted experiments. K.M.L. provided conceptual insights and assisted experiments. W.J.G. advised on data analysis and provided conceptual insights. All authors read and approved of the manuscript.

## Experimental Procedures

### Cell lines

FaDu, a pharyngeal squamous cell carcinoma cell line, was purchased from ATCC (HTB-43) and cultured in ATCC-formulated Eagle’s Minimum Essential Medium supplemented with 10% fetal bovine serum (FBS; Omega Scientific) and antibiotics (100 units/mL Penicillin and 100 µg/mL Streptomycin; Gibco). NCI-H520 lung squamous cell carcinoma cells were purchase from ATCC (HTB-182) and cultured in RPMI-1640 medium (ATCC modification, Gibco) supplemented with 10% FBS and antibiotics. T.T esophageal squamous cell carcinoma cells were purchase from JCRB Cell Bank (JCRB0262) and cultured in medium Dulbecco’s Modified Eagle Medium: Nutrient Mixture F-12 (DMEM/F-12; Gibco, catalog no. 10565) supplemented with 10% FBS, and antibiotics. KYSE70 esophageal squamous cell carcinoma cell line was purchase from Sigma (94072012) and cultured in medium RPMI-1640 (Gibco, catalog no. 21870092) supplemented with 10% FBS, 10 mM HEPES (Gibco), 1 mM sodium pyruvate (Gibco), 2mM GlutaMax (Gibco) and antibiotics. Primary normal human epidermal keratinocytes (KC) were purchased from Gibco (C0055C) and cultured on Geltrex (Gibco) coated plates with EpiLife basal medium with 60 µM calcium (Gibco) plus 100 µg/mL insulin, 15 µg/mL transferrin, 10ng/mL epidermal growth factor, 15 µM forskolin, 500nM VX-745, 250nM RO4929097, 100nM dexamethasone and antibiotics. HEK293T cells were purchased from Takara Bio USA (632180) and cultured in high glucose DMEM (GIBCO) medium supplemented with 10% FBS (GIBCO), 10 mM HEPES (Gibco), 1 mM sodium pyruvate (Gibco), 2mM GlutaMax (Gibco) and antibiotics. Cells were maintained in a humidified incubator at 37 °C in the presence of 5% CO_2_ and passaged every 2–3 days. Cell lines were routinely tested for mycoplasma and immediately tested upon suspicion. None of the cell lines used in the reported experiments tested positive.

### Estimate of protein molecules

For the preparation of the whole-cell lysates, the cells were lysed in RIPA buffer (50 mM Tris-HCl pH 8.0, 300 mM NaCl, 1 mM EDTA, 0.1% SDS, 1% NP-40, 0.5% sodium deoxycholate, 20 mM NaF, 1 mM DTT, 1 mM sodium orthovanadate, 0.25 mM PMSF and protease inhibitors) supplemented with 5 mM MgCl_2_ and 0.5 U/ul of Benzonase (Sigma). After the samples were sat on ice for 30 minutes, the LDS sample buffer with final 2.5% β-mercaptoethanol was added, followed by boiling for 5 minutes. The extracts from 300,000 cells were subjected to SDS–PAGE and western blot analysis together with 1.25 ng, 5 ng, 10 ng, and 20 ng of purified ACTL6A or SMARCA4 recombinant proteins. Odyssey CLx LI-COR was used to analyzed and quantified the Western blot signals. The standard curves of signal to mass from the recombinant proteins were applied for estimating the amount of ACTL6A or SMARCA4/SMARCA2 from the cell lysates, which was further divided by the cell number (300,000) to obtain the mass (g) per cell, and then the number of molecules (N) per cell calculated by the following formula: *N= mass (g) / (molecular weight (kDa) x10^3^) x Avogadro constant (6.022x10^23^)*

For producing recombinant proteins, the DNA fragment encoding human ACTL6A amino acid 43-119, the region used to raise the anti- ACTL6A antibodies (Crabtree laboratory), was inserted between the BamHI and HindIII sites of pGSTag (Addgene 21877); and the fragment expressing human SMARCA4 amino acid 1086-1307, used to raise the J1 antibodies (Crabtree laboratory) that recognize both SMARCA4 and SMARCA2, was inserted between the BamHI and HindIII sites of pMAL-c2X (Addgene 75286). After 0.4mM IPTG induction for 2 hours at 37°C, the bacteria were collected and resuspended in PBS with 1mM EDTA, 1mM PMSF, 0.02% (∼3mM) β-mercaptoethanol, and protease inhibitors. The cells were lysed by Diagenode Bioruptor for 15 min, high output. After 3,500 rpm spin for 10 mins, the supernatants were rotated with amylose resin (New England Biolabs) for MBP tag, or glutathione-superflow resin (Clontech) for GST tag overnight at 4°C. The resins were washed four times by PBS supplemented with 350mM NaCl, 0.1% triton-X-100, 1mM EDTA, 1mM PMSF, 0.02% (∼3mM) β-mercaptoethanol, and protease inhibitors. The GST-tagged proteins were eluted by 10mM reduced glutathione (Sigma, 100mM stock made in 50mM Tris, pH7.6) in PBS (containing 1mM PMSF and 0.02% (∼3mM) β-mercaptoethanol), and MBP-tagged proteins were eluted by 10mM maltose.

### ACTL6A knockdown

siRNA transfections were performed using DharmaFECT 1 transfection reagents (Horizon Discovery) in the antibiotics-free medium according to the manufacturer’s instructions. The siRNA reagents were purchased from Horizon Discovery (ON-TARGETplus Human ACTL6A siRNA: Dharmacon L-008243-00; control ON-TARGETplus Non-targeting Pool: Dharmacon D-001810-10) and resuspended in siRNA buffer (Horizon Discovery). CRISPR gRNAs were cloned into vector lentiCRISPR v2 (Addgene 52961). The sequences of human *ACTL6A* targeting gRNAs are: TAATGCTCTGCGTGTTCCGA, ATGAGCGGCGGCGTGTACGG, GCGTGTTCCGAGGGAGAATA, AGATGACGGA-AGCACATTAA.

For producing lentiviral particles, lentiviral vectors (18 μg per 15 cm^2^ dish) together with packaging vectors pMD2.G (4.5 μg) and psPAX2 (13.5 μg) were delivered into lenti-X 293T cells (Clontech) using 108-144 μg PEI MAX 40K (Polysciences, cat. 24765; stock 1 μg /ul) mixed in 1.8 ml Opti-MEM (Gibco) according to the manufacturer’s instructions. 12-16 hours after transfection, the medium was replaced by viral production medium (UltraCULTURETM serum-free cell culture medium (Lonza) supplemented with 10mM HEPES (Gibco), 1 mM sodium pyruvate (Gibco), 2mM GlutaMax (Gibco) and antibiotics (Gibco)). 72 hr post-transfection, lentiviral particles were collected by centrifugation of 0.45 μm pore size-filtered cell culture supernatants at 20,000 rpm for 2 hours at 4 °C, followed by PBS resuspension. Lentiviral transduction was conducted by Spinfection method in the presence of 10 μg/ml polybrene at 1,100 x*g* for 30 min at 37°C. 48 hours post-infection, infected cells were selected by 2μg/ml puromycin (Sigma), followed by 10μg/ml Blasticidin (Gibco) in *ACTL6A* reconstitution experiments.

### ACTL6A over-expression

Human *ACTL6A* cDNA were cloned between Not1 and Mlu1 restriction enzyme sites downstream of the EF1α promoter in Crabtree lentiviral vector N103 (puromycin selection) and N106 (blasticidin selection), which harbor a second promoter PGK driving drug resistance gene for selection. See the “ACTL6A knockdown” section for lentiviral production and infection. For site-directed mutagenesis, *ACTL6A* cDNA were cloned into pUC-19 vector between HindIII and Kpn1, and the construct was used as template in PCR with primers- for P373S/P374G: TCAGAAAACTTCTGGAAGTATGCGG, GACAGCTCTCTATTCAAC; for R377G: TCCAAGTATGGGCTTGAAATTGATTGC, GGAGTTTTCTGAGACAGC. The PCR products were treated by kinase, ligase and Dpn1 (KLD) enzyme mix (New England Biolabs), followed by bacterial transformation, clone selection by Sanger DNA sequencing, and subcloning back to lentiviral vector N103 (puromycin selection) and N106 (blasticidin selection) between Not1 and Mlu1 sites. For generating *ACTL6A* expressing constructs that are resistant to *ACTL6A*-CRISPR KO, following silent mutations marked by underlines were introduced to block *ACTL6A* gRNA binding: atg<u>TCT</u>gg<u>A</u>gg<u>A</u>gt<u>C</u>ta<u>T</u>gg, <u>G</u>ga<u>C</u>ga<u>T</u>gg<u>CTCT</u>ac<u>C</u>tt<u>G</u>a, <u>C</u>aa<u>C</u>gc<u>C</u>ct<u>CAGG</u>gt<u>C</u>cc-<u>TC</u>g<u>C</u>ga<u>A</u>aata.

### Immunoprecipitation and Western blot

For protein-protein interaction studies, cells reaching 80-90% confluence on the culture plates were washed once by cold PBS and lysed in cold hypotonic lysis buffer A, ∼0.5ml/10cm^2^ growth area (Buffer A: 25 mM HEPES pH 7.5, 25 mM KCl, 0.05 mM EDTA, 5mM MgCl_2_, 10% glycerol, 0.1% NP-40, 1 mM DTT, 1 mM sodium orthovanadate, 0.25 mM PMSF and protease inhibitors). After incubated on ice for 5 minutes, cells were scraped from plates by cell lifter, harvested, and spun down at 1500 rpm for 5 mins at 4 °C. Then, the nuclei were washed by buffer A twice. For each wash, cold 5ml buffer A per 20-million cells were added to the pellet, followed by 5-minute incubation on ice, centrifugation at 1,500 rpm for 5 mins, and discarding the supernatants. The nuclei were lysed in immunoprecipitation (IP) buffer, ∼1 ml per 20-million cells (IP buffer: 20 mM HEPES pH7.5, 100 mM KCl, 2.5 mM MgCl_2_, 5% glycerol, 1% Triton X-100, 0.5% NP-40, 1 mM dithiothreitol (DTT), 1mM sodium orthovanadate, 0.25 mM phenylmethylsulphonylfluoride (PMSF) and protease inhibitors). The nuclei were further passed through a 1 ml 27G-needle syringe 5 times and sonicated for three cycles of 10 sec-ON/1 minute-OFF, high output (Diagenode Bioruptor). After centrifugation at 15,000 rpm for 10 min to collect the supernatants, the concentrations of the nuclear extracts were measure by Bradford assay (Bio-Rad) and adjusted to 1-1.5 µg/µl by IP buffer. For each IP, 500-1000ug protein extracts were incubated under rotary agitation overnight at 4 °C with antibodies against SMARCA4 (Santa Cruz Biotechnology, sc-374197 X; 4 µg), YAP (Cell Signaling Technology, 14074S; 5 µl), Pan-TEAD (Cell Signaling Technology, 13295S; 5 µl), TEAD4 (Abcam, ab58310; 3ug), ACTL6A (Invitrogen, 702414; 2.5 µg), or mouse/rabbit IgG control (Santa Cruz Biotechnology sc-2025; MilliporeSigma, 12-370). After additional one hour incubation with 40ul Protein A or G dynabeads (Invitrogen), the beads were washed four times by 1 ml IP buffer and resuspended in 20 µl 1x LDS sample buffer (Invitrogen) containing 2.5% β-mercaptoethanol and then boiled at 95 °C for 5 min. The samples were subjected to SDS–PAGE and western blot analysis. For *ACTL6A* perturbations, cells were collected 72 hours after siRNA transfection or 5 days after infection by lentiCRISPR, or 1-2 weeks after infection by lentivirus carrying *ACTL6A*.

Antibodies used for Western blotting included those against SMARCC1 (Crabtree laboratory), SMARCA4 (Santa Cruz Biotechnology, sc-374197 X), ACTL6A (Novus Biologicals, NB100-61628; or homemade in the Crabtree laboratory), J1 SMARCA4/SMARCA2 (Crabtree laboratory), ARID1A (Santa Cruz Biotechnology, sc-32761), BAF57 (Bethyl Laboratories, A300-810A), YAP (Abnova, H00010413-M01; Cell Signaling Technology, 14074S), Phospho-YAP Ser127 (Cell Signaling Technology, 13008), Pan-TEAD (Cell Signaling Technology, 13295S), TEAD1 (BD Biosciences, 610922), TEAD4 (Abcam, ab58310), INO80 (Bethyl Laboratories, A303-371A), EZH2 (BD Biosciences, 612666), GAPDH (Cell Signaling Technology, 5174S), V5 (Invitrogen, R960-25).

### Density gradient sedimentation analysis

A detailed description has been published elsewhere (Lessard et al., 2007). In brief, 30-40 million cells were washed by PBS once and lysed in 10 ml cold hypotonic lysis buffer A (see Immunoprecipitation) and then sat on ice for 7 minutes. After centrifugation at 1500 rpm for 5 mins at 4 °C, nuclei were further washed by 10 ml buffer A twice and re-suspended in 700 µl buffer C (10 mM HEPES, pH7.5, 100 mM KCl, 0.1 mM EDTA, 3mM MgCl_2_, 10% glycerol, 1 mM DTT, 1mM sodium orthovanadate, 0.25 mM PMSF and protease inhibitors). Chromatin proteins were extracted with 0.3M ammonium sulfate (pH 7) by adding 1/9 volume of 3M ammonium sulfate stock and incubated under rotary agitation for 1-2 hours at 4 °C. Nuclear extracts were collected after ultracentrifugation at 100,000 rpm for 15 minutes at 4 °C (TLA 120.2 rotor), and proteins were precipitated with 0.33 mg/µl ammonium sulfate on ice for 20 min. Precipitated proteins were pelleted by another ultracentrifugation at 100,000 rpm for 15 minutes at 4 °C and re-suspended in 200 µl HEMG-0 buffer (25 mM HEPES, pH7.9, 100 mM KCl, 0.1 mM EDTA, 12.5mM MgCl_2_ supplemented with 1 mM DTT, 1mM sodium orthovanadate, 0.25 mM PMSF and protease inhibitors). Protein concentration was measured by Bradford assay (Bio-Rad) and adjusted accordingly for glycerol gradient analyses. 200 µl of the solution with 500-1000 µg proteins were overlaid on a 10-ml density-gradient liquid column with 10 to 30% glycerol (in HEMG buffer) and placed in a SW-40 swing bucket rotor for centrifugation at 40,000 rpm for 16 h at 4 °C. A series of 0.5ml fractions were then recovered from the top, and further subject to SDS–PAGE and western blot analysis.

### Subcellular fractionation

Cells were first lysed in cold hypotonic lysis buffer A (Buffer A: 25 mM HEPES pH 7.5, 25 mM KCl, 0.05 mM EDTA, 5mM MgCl_2_, 10% glycerol, 0.1% NP-40, 1 mM DTT, 1 mM sodium orthovanadate, 0.25 mM PMSF and protease inhibitors). After incubated on ice for 5 minutes, cells were spun down at 1500 rpm for 5 mins at 4 °C. The supernatants were collected as cytoplasmic fraction. Then, the nuclei were washed by buffer A twice and lysed in equal volume to the cytoplasmic fractions of RIPA buffer (50 mM Tris-HCl pH 8.0, 300 mM NaCl, 1 mM EDTA, 0.1% SDS, 1% NP-40, 0.5% sodium deoxycholate, 20 mM NaF, 1 mM DTT, 1 mM sodium orthovanadate, 0.25 mM PMSF and protease inhibitors) supplemented with 5 mM MgCl_2_ and 0.5 U/ul of Benzonase (Sigma); and then sit on ice for 30 minutes. Both cytoplasmic fractions and nuclei solutions were subject to centrifugation at 15,000 rpm for 10 min at 4 °C to remove the cell debris. The samples were added with LDS sample buffer (Invitrogen)/β-mercaptoethanol, boiled for 5 minutes, and analyzed by Western blot.

### Immunofluorescence

24 hours after siRNA transfection, cells were re-seeded on chamber slides coated by fibronectin (coating: 20 µg/ml fibronectin (Sigma) at 37 °C overnight). 72 hours post-transfection, cells were fixed in 4% paraformaldehyde (PFA) for 10 min at room temperature (RT). For immunostaining, cells were permeabilized in PBS with 0.3% Triton X-100 for 20 min and blocked for 1 h at RT in blocking buffer (PBS containing 2.5 % normal donkey serum, 2.5 % normal goat serum, 1% BSA and 0.1% Triton X-100) supplemented with M.O.M. blocking reagent (Vector Laboratories). Primary YAP (Abnova, H00010413-M01) were diluted in blocking buffer supplemented with M.O.M. Protein Concentrate (Vector Laboratories) and applied to cells, followed by overnight incubation at 4 °C. After washing by PBS with 0.1% Triton X-100 three times at RT, cells were incubated for 1 hour at RT with secondary antibodies conjugated to Alexa-488 (Invitrogen). The slides were mounted with ProLong Diamond Antifade Mountant with DAPI (Invitrogen). Images was captured on the Keyence BZ-X700 microscope, and prepared by ImageJ, Adobe Photoshop, and Illustrator CS6.

### RNA extraction, RT-qPCR, and RNA-seq analysis

72 hours after siRNA transfection or one week after lentiviral transduction and drug selection, cells were lysed directly on culture plates with TRIsure reagents (Bioline), and RNA was extracted using Direct-zol RNA MiniPrep kits (Zymo Research) with in-column DNase I digestion to remove residual genomic DNA. For RT-qPCR, complementary DNA was further synthesized using SensiFAST cDNA synthesis kit (Bioline). cDNAs were mixed with indicated primers and SensiFAST SYBR lo-ROX reagents (Bioline), and quantitative PCR (qPCR) was performed on Applied Biosystems QuantStudio 6 Flex Real-Time PCR System. Specificity was confirmed by subsequent melting curve analysis or gel electrophoresis. Levels of PCR products were expressed as a function of peptidylprolyl isomerase B (*PPIB*). Primers were designed through Primer 3 or from previous reports, and amplified products encompass exon/intron boundaries. The sequences of primers used in this study:

*<u>ACTL6A</u>* forward 5’- AGTGGCAGGAGGAAACACAC -3’, reverse 5’-CCCAAAGAGGCTAGAATGGA -3’;

*<u>CTGF</u>* forward 5’- AGGAGTGGGTGTGTGACGA -3’, reverse 5’- CCAGGCAGTTGGCTCTAATC -3’;

*<u>AXL</u>* forward 5’-CACCAGCAAGAGCGATGTGT -3’, reverse 5’- CGGTCCTGGGGATTTAGCTC -3’;

*<u>OLR1</u>* forward 5’- TTGCCTGGGATTAGTAGTGACC -3’, reverse 5’- GCTTGCTCTTGTGTTAGGAGGT -3’;

*<u>TGM2</u>* forward 5’- GGCACCAAGTACCTGCTCA -3’, reverse 5’- AGAGGATGCAAAGAGGAACG -3’;

*<u>BRPF3</u>* forward 5’- GGAAGGTGTGAACGGAGACT -3’, reverse 5’- GTCTCCGCGGTCTTCAAA -3’;

*<u>CAVIN2</u>* forward 5’- GGCAGGTATGACAGCTTACGG -3’, reverse 5’- GTTGTCCACCGGCTGTAATGT -3’;

*<u>IGFBP3</u>* forward 5’- AGAGCACAGATACCCAGAACT -3’, reverse 5’- GGTGATTCAGTGTGTCTTCCATT -3’;

*<u>FNDC3B</u>* forward 5’- GAGCATGCTGCATCAGTACC -3’, reverse 5’- GTGCGAACAGGGAGACTTTC -3’;

*<u>TYRO3</u>* forward 5’- GAGAGGAACTACGAAGATCGGG -3’, reverse 5’- AGTGCTTGAAGGTGAACAGTG -3’;

*<u>CNN3</u>* forward 5’- GGCAGGTATGACAGCTTACGG -3’, reverse 5’- GTTGTCCACCGGCTGTAATGT -3’;

*<u>WWC1</u>* forward 5’- TCTACCAGGTGAAGCAGCAG -3’, reverse 5’- GCTGGATGATGAACCAGAGAC -3’;

*<u>PPIB</u>* forward 5’- CGTCTTCTTCCTGCTGCTG -3’, reverse 5’- AGCCAAATCCTTTCTCTCCTG -3’.

RNA-seq libraries were generated by using NEBNext ultra II directional RNA library prep kit coupled with NEBNext multiplex oligos for Illumina (New England Biolabs) and following the manufacturer’s directions. The deep sequencing was performed on the NextSeq 550 sequencing system (Illumina) with a 75-cycle, paired-end sequencing. Alignment of RNA-sequencing reads was performed with STAR (Dobin et al., 2013) with ENCODE standard options to GENCODE v19, and read counts were generated using eXpress (Roberts and Pachter, 2013). Differential gene expression was determined using DEseq2 (Love et al., 2014).

### CUT&RUN and data analysis

CUT&RUN was performed as previously described (Skene et al., 2018). Cells were collected 72 hours after siRNA transfection or one week after lentiviral transduction and drug selection. 250,000 cells were bound to 10 µl concanavalin-A beads and collected into a 0.2 ml PCR tube. Antibody binding was conducted in the volume of 50 µl on thermomixer for 96-well PCR plates (Eppendorf) in 4 °C cold room for 2 hours, and Protein A-MNase binding for 1 hour. Wash buffer volume was adjusted to 150 µl. The digestion was performed under high Ca^2+^/low salt condition on ice for 15 minutes, followed by incubation at 37 °C for 30 min to release CUT&RUN fragments. Antibodies were diluted 1/100 and included: YAP (Cell signaling, 14074T), TEAD1 (Cell signaling, 12292S), SMARCC1 (Crabtree laboratory), H3K4me1 (Abcam, ab8895), H3K4me3 (Active motif, 39159), H3K27Ac (Abcam, ab4729), H3K27me3 (cell signaling, 9733). For histone marks profiling, CUT&RUN fragments were purified by spin column-based methods (Zymo DNA clean & concentrator-5 kit). For transcription factors and BAF complex profiling, CUT&RUN fragments were extracted by phenol-chloroform to recover small fragments.

The libraries were generated by using the NEBNext ultraII DNA library prep kit for Illumina coupled with NEBNext multiplex oligos for Illumina (New England Biolabs) with few modifications optimized for small fragments (detailed in <u>dx.doi.org/10.17504/protocols.io.wvgfe3w</u>.) The deep sequencing was performed on the NextSeq 550 sequencing system (Illumina) with 75-cycle, paired-end sequencing setup. Reads were first trimmed from Illumina adapters using SeqPurge (Sturm et al., 2016). Demultiplexed fastq files were mapped to the hg19 assembly of the human genome as 2×36 mers using Bowtie2 (Langmead and Salzberg, 2012), and generated BAM files were then filtered for low-quality reads and duplicated reads. TEAD1 and YAP mapped fragments <120bp were used for downstream analysis. Coverage bigwigs were generated using bamCoverage from deeptools (Ramirez et al., 2016) with CPM normalization. Peaks were called using macs2 (Zhang et al., 2008) with the nomodel option. Fixed-width peaks from replicates and treatments were combined using the called peaks summits files as described previously (Corces et al., 2018). *De novo* motif search on all CUT&RUN libraries was done using Homer (v4.11) (Heinz et al., 2010) with default parameters. Counts over peaks were generated using the featureCounts (Liao et al., 2014).

Differential peaks were determined using DEseq2 (Love et al., 2014) or edgeR (Robinson et al., 2010). To annotate joined YAP/TEAD1/SMARCC1 peaks, we defined the following states based on CUT&RUN datasets: active promoter (high H3K4me3 & low H3K27me3 & TSS distance < 1000), poised promoter (high H3K4me3 & high H3K27me3 & TSS distance < 1000), active enhancer (high H3K4me1 & high H3K27Ac & TSS distance > 1000), poised enhancer (high H3K4me1 & high H3K27me3 & TSS distance > 1000) and repressed (H3K27me3 high). High and low were defined by 0.99 percentile. Differential analysis for H3K27me3 CUT&RUN between *ACTL6A* -overexpressing and vector-control keratinocytes was performed using DiffBind (Ross-Innes et al., 2012) by binning the genome into 5- kb bins and performing differential analysis (using DESeq2) between two conditions. We identified 4,035 H3K27me3-differential bins (constituting 2389 broad peaks), of which 1,963 showed decreased H3K27me3 levels upon *ACTL6A* over-expression. Genes with H3K27me3 differentials in their promoters were defined as genes whose TSS is within the differential H3K27me3 5-kb bin. H3K27me3 CUT&RUN and H3K4me3 ChIP-seq (ENCSR075OQB for keratinocytes from ENCODE (Zhang et al., 2020)) signals around gene promoter with differential H3K27me3 levels was performed using computeMatrix and plotProfile from deeptools (Ramirez et al., 2016). H3K27me3 ChIP-seq for keratinocytes were from ENCODE ENCSR377MRR (Zhang et al., 2020). Heatmaps for CUT&RUN and ATAC-seq data around different genomic featutres were done using deeptools (v2.5.6) (Ramirez et al., 2016) computeMatrix. Plots were done using deeptools plotHeatmap or plotProfile option. Differential gene expression data of RNA-seq for lung squamous cell carcinoma (LUSC) and head-and-neck squamous cell carcinoma (HNSC) versus their normal tissues were from GEPIA2 (Tang et al., 2017) with cutoff *p*-value < 0.05. HNSC and LUSC datasets were merged by gene name and then merged with genes displaying differential H3K27me3 CUT&RUN signals to identify ACTL6A-dependent PRC target genes that were preferentially altered in either HNSC or LUSC tumors.

### Omni-ATAC-seq and data analysis

We followed the Omni-ATAC method in Corces MR *et al*. (Corces et al., 2017). 72 hours after siRNA transfection or one week after lentiviral transduction and drug selection, cells were pretreated with 200 U/ml DNase (Worthington) for 30 min at 37 °C to remove free-floating DNA and DNA from dead cells. Nuclei from 75,000 cells were used in the transposition reactions (Illumina) at 37 °C for 30 min, followed by DNA purification by Zymo DNA clean & concentrator-5 kits, and library preparation. Deep sequencing was conducted on the NextSeq 550 sequencing system (Illumina) with 75- cycle, paired-end sequencing. Demultiplexed and trimmed fastq files were mapped to the hg19 assembly of the human genome as 2×36 mers using Bowtie2 (Langmead and Salzberg, 2012). Duplicate reads were removed using picard-tools (v.1.99). Low-quality reads and chrM reads were removed using samtools. Fixed width peaks were called using macs2 (Zhang et al., 2008) with the nomodel option, and summits were called. Peaks from replicates and treatments were combined using the called peaks summits files as described previously (Corces et al., 2018). Read counts over the peaks were done using the ChrAccR package (https://github.com/GreenleafLab/ChrAccR). Differential peaks were determined using DEseq2 (Love et al., 2014) or edgeR (Robinson et al., 2010).

To find transcription factors that are enriched in differential peaks, we used motifMatcher (part of chromVar (Schep et al., 2017) package to call TF binding sites within the differential peak sets. Using hypergeometric test, we computed the enrichment of each TF within increasing/decreasing differential peaks with all peaks as background. Heatmaps for CUT&RUN and ATAC-seq data around different genomic featutres were done using deeptools (Ramirez et al., 2016) (v2.5.6) computeMatrix. Plots were done using deeptools plotHeatmap or plotProfile option. Genomic feature annotation for ATAC-seq peaks that gained or lost accessibility as a result of *ACTL6A* loss was performed using ChiPseeker (Yu et al., 2015b).

### Statistics

Comparisons were performed in Prism 8 (GraphPad Software) with unpaired two-tailed student’s t-test to determine significance between two groups indicated in figures. The cancer genomic data were from cBioPortal TCGA PanCancer Atlas studies (Cerami et al., 2012; Gao et al., 2013), and the RNA analyses of tumors versus paired normal tissues were from GEPIA 2 (Tang et al., 2017).

### Data availability

Next-generation sequencing data reported in this study have been deposited at the Gene Expression Omnibus with accession number GSE156788. Protein structures for BAF complex were from Protein Data Bank (https://www.rcsb.org, PDB IDs: 6LTJ).

### Code availability

All codes for analyses and generating plots herein will be available upon request.

