## Supplemental figures for "Increased ACTL6A Occupancy Within mSWI/SNF Chromatin Remodelers Drives Human Squamous Cell Carcinoma"

**This PDF file includes:**

Figures S1 to S5

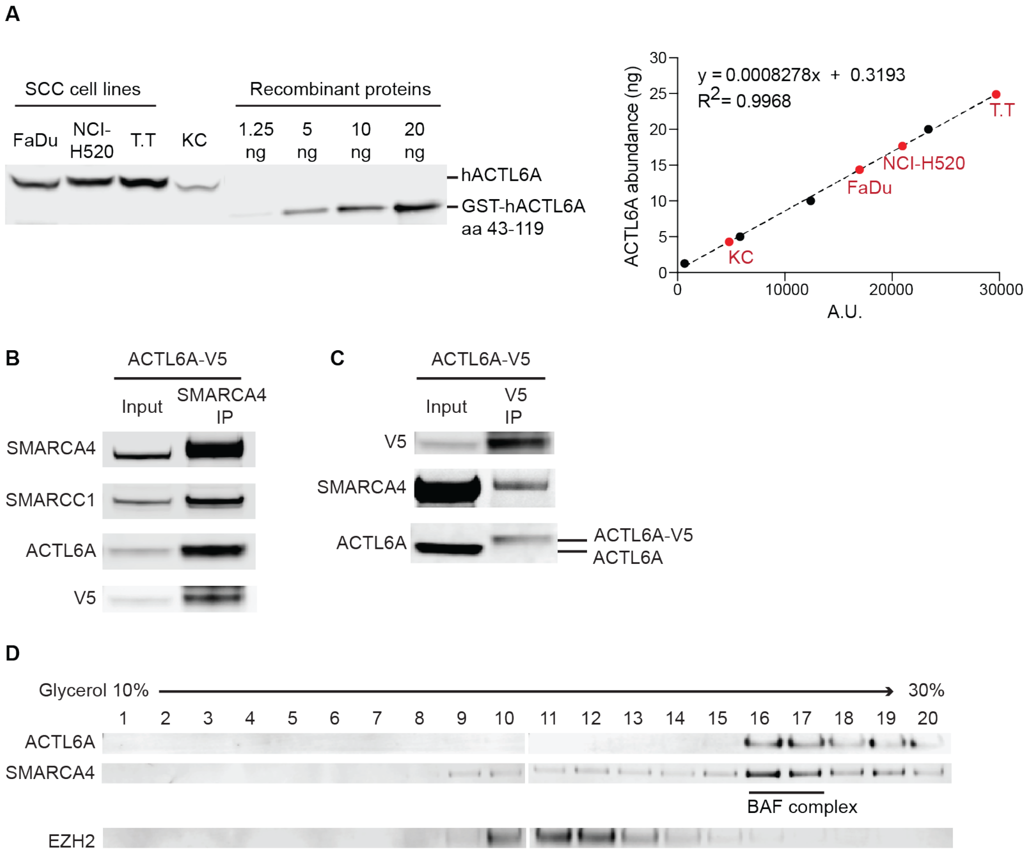


**Figure S1.** ***ACTL6A*** **over-expression does not induce its dimerization and most of ACTL6A proteins associate with BAF complexes in SCC cells. Related to Figure 2.**
(A) Left: Western blots for whole cell lysates from 300,000 cells of indicated cell lines and ACTL6A recombinant proteins. SCC cell lines: FaDu (head-and-neck), NCI-H520 (lung), and T.T (esophageal). KC: primary normal human keratinocytes. Right: Quantifications for Western blots. A.U.: arbitrary unit of Western blot integrated intensity. Linear regression line was generated from serially diluted ACTL6A recombinant proteins (black circles) and used for calculating ACTL6A abundance (ng) in cells (red circles). (B) Co-IP by SMARCA4 antibody using nuclear extracts from FaDu SCC cells infected with lentivirus expressing C-terminal V5-tagged ACTL6A. Note the incorporation of ACTL6A-V5 into BAF complexes as endogenous ACTL6A. (C) Co-IP by V5 antibody as (B). Note endogenous ACTL6A does not bind V5-tagged ACTL6A. (D) Density sedimentation and immunoblots for nuclear extracts from SCC cells. Note ACTL6A co-migrated with SMARCA4 marking BAF complexes in high molecular mass fractions 16-17 as a full complex. EZH2: subunit of PRC2 complexes as a control.

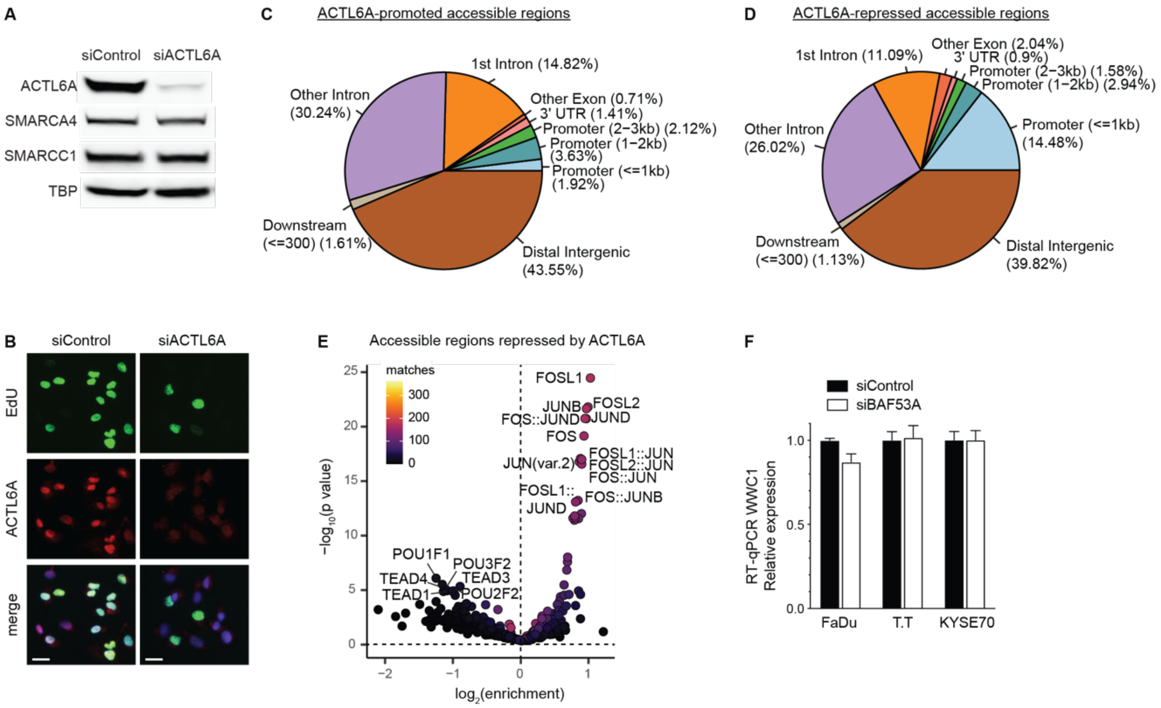


**Figure S2. ATAC-seq analyses identify accessible chromatin regions regulated by ACTL6A in SCC cells. Related to Figure 3.**
(A) Western blots for cell lysates from FaDu SCC cells 72h after transfection with *ACTL6A* siRNA (siACTL6A) or control siRNA (siControl). TBP: TATA-box-binding protein as loading control. (B) Immunostaining showing decreased SCC cell proliferation in response to *ACTL6A* knockdown. Assessment by the incorporation of EdU (5-ethynyl-2'-deoxyuridine) applied 1.5h before analysis, 72 hours after siRNA transfection. Scale bars: 20 μm. (C) Annotation pie chart of regions with decreased accessibility by siACTL6A compared to siControl in FaDu SCC cells from ATAC-seq analysis. (D) As in (C) for regions with increased accessibility by siACTL6A compared to siControl. (E) Transcription factor (TF) motif enrichment within regions with increased accessibility in siACTL6A versus siControl FaDu cells. Matches: number of peaks containing matched TF binding motifs. (F) RT-qPCR showed no significant changes in *WWC1* expression levels 72 hours after siACTL6A transfection compared to siControl in three SCC cell lines. Mean ± SEM. *$P < 0.05$.

A

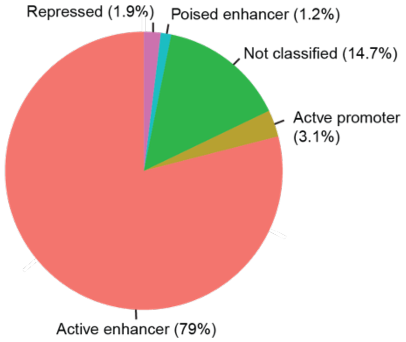


B

6,251 joint peaks between YAP, TEAD1 and SMARCC1 CUT&RUN

| Rank | Motif | P-value | % of Targets | Best Match |
|---|---|---|---|---|
| 1 | 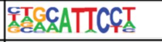 | $1x10^{-1146}$ | 64.49% | TEAD4 (TEA) |
| 2 | 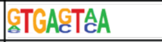 | $1x10^{-265}$ | 55.22% | Fos (bZIP) |
| 3 | 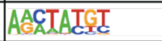 | $1x10^{-58}$ | 28.08% | NEUROG2 |
| 4 | 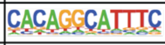 | $1x10^{-39}$ | 11.45% | POL009.1_DCE_S_II |
| 5 | 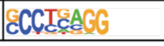 | $1x10^{-30}$ | 14.33% | TFAP2E |

C

40,551 peaks from SMARCC1 CUT&RUN

| Rank | Motif | P-value | % of Targets | Best Match |
|---|---|---|---|---|
| 1 | 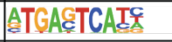 | $1x10^{-5642}$ | 56.09% | FOSL2::JUNB |
| 2 | 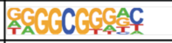 | $1x10^{-444}$ | 6.79% | Sp2(Zf) |
| 3 | 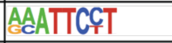 | $1x10^{-126}$ | 20.94% | TEAD(TEA) |
| 4 | 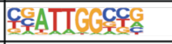 | $1x10^{-123}$ | 6.59% | NFY(CCAAT) |
| 5 | 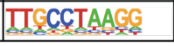 | $1x10^{-112}$ | 46.26% | TFAP2A |

D

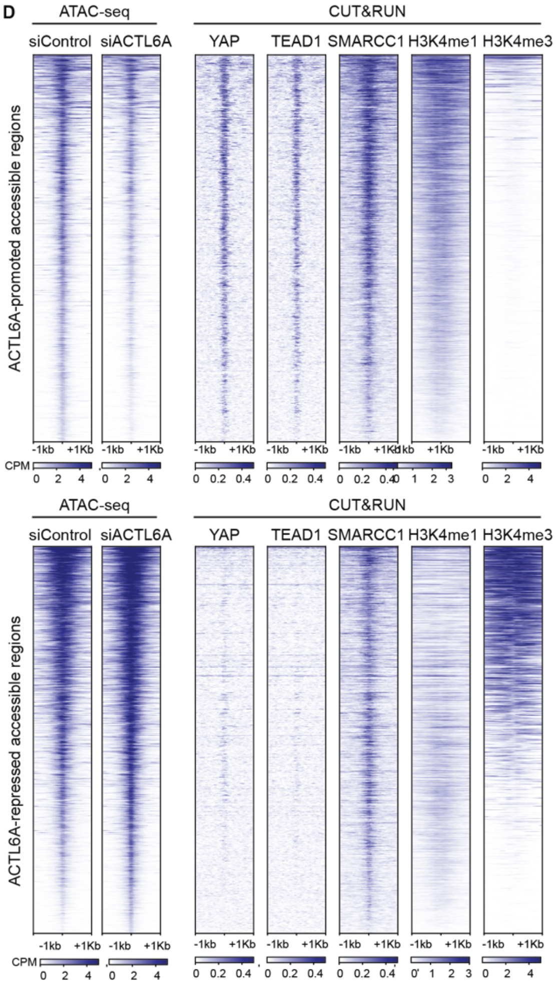


E

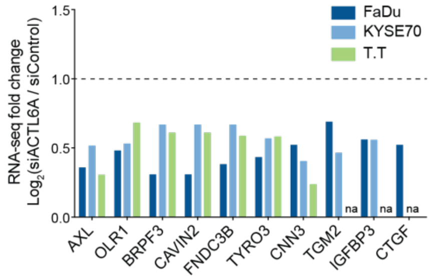


F

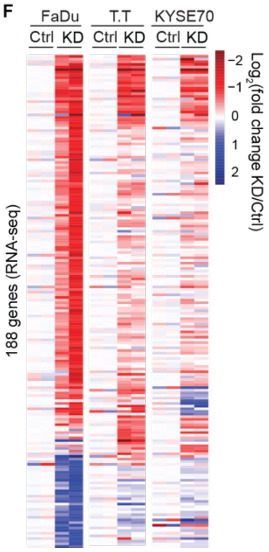


G

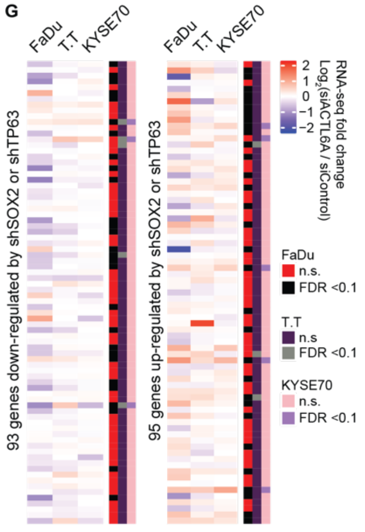

**Figure S3. CUT&RUN profiling reveals the co-localization of YAP, TEAD1 and BAF complexes on chromatin in SCC cells. Related to Figure 3.**
(A) Distribution of YAP, TEAD1 and BAF-subunit SMARCC1 co-occupied regions identified by CUT&RUN in FaDu SCC cells. Classification is based on the presence of histone marks identified by CUT&RUN: active enhancers- H3K27Ac$^{+}$ H3K4me1$^{+}$ (79 %); poised enhancers- H3K4me1$^{+}$ H3K27me3$^{+}$ (1.2 %); active promoters- H3K4me3+ H3K27me3$^{-}$ (3.1 %); poised promoter- H3K4me3+ H3K27me3$^{+}$ (0 %); repressed- H3K27me3$^{+}$(1.9 %). (B) *De novo* motif analysis by HOMER for regions co-bound by YAP, TEAD1 and SMARCC1 identified by CUT&RUN. (C) As in (B), for regions bound by SMARCC1. (D) Heat maps for ATAC-seq and indicated CUT&RUN profiles around regions with decreased accessibility (top) and increased accessibility (bottom) in siACTL6A relative to siControl FaDu SCC cells. H3K4me1: enhancer histone mark. H3K4me3: active promoter histone mark. (E) Relative expression levels of YAP target genes from RNA-seq analyses in siACTL6A cells normalized to siControl values. na: not available (not expressed). (F) Hierarchical clustering of RNA-seq analyses showing 188 genes with differential expression between siACTL6A (KD) and siControl (Ctrl) in at least two of three SCC cell lines. FDR < 0.05. (G) Heat maps showing the RNA level fold-changes between siACTL6A versus siControl in three SCC cell lines across SOX2 and TP63 co-regulated genes. *SOX2* shRNA (shSOX2) or *TP63* shRNA (shTP63) knockdown experiments (Watanabe et al., 2014) were done in KYSE70 SCC cell line. Binary code on the right: FDR< 0.1 or not significant (n.s.).

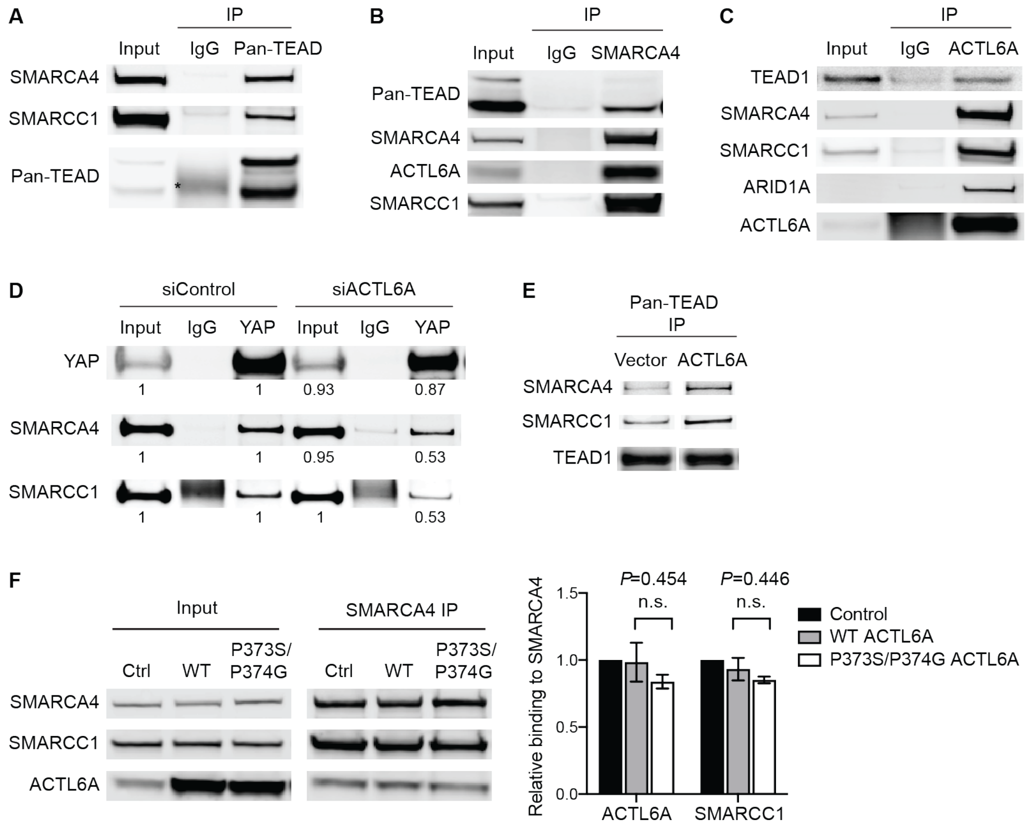


**Figure S4. ACTL6A provides surfaces on the BAF complex for TEAD-YAP binding. Related to Figure 4.**

(A-C) Co-IP experiments by Pan-TEAD (A), SMARCA4 (B), ACTL6A (C) antibodies with control IgG using nuclear extracts from FaDu SCC cells. *IgG heavy chain. (D) Co-IP by YAP and control IgG antibodies using nuclear extracts from FaDu cells 72 hours after transfection with siACTL6A and siControl. Relative levels normalized to siControl condition. (E) Co-IP experiments by Pan-TEAD antibody in primary human keratinocytes transduced by lentivirus overexpressing *ACTL6A* and the vector control. Note increased binding of BAF subunit SMARCA4 and SMARCC1 to TEAD. Mouse TEAD1 antibody used for blotting to avoid TEAD signals masked by rabbit Pan-TEAD antibody heavy chain signals. (F) Co-IP by SMARCA4 antibody in nuclear extract from FaDu cells. WT: reconstituting WT ACTL6A into *ACTL6A* CRISPR-KO cells. P373S/P374G: reconstituting P373S/P374G ACTL6A into *ACTL6A* CRISPR-KO cells. Ctrl: lentiviral CRISPR control vector. Quantifications: relative levels of co-IP'd ACTL6A and SMARCC1 in WT and P373S/P374G conditions normalized to ctrl. n=2 experiments. Error bars indicate SEM. n.s.: not significant.

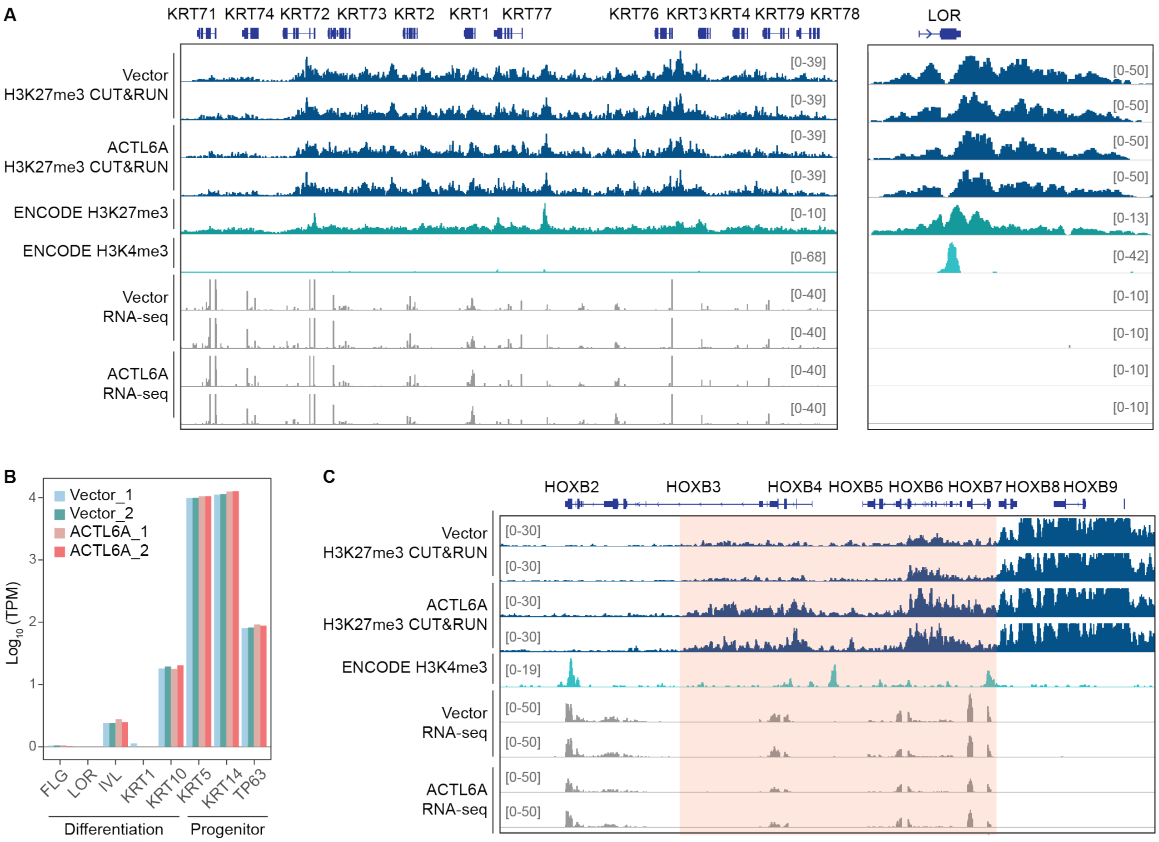


**Figure S5. Polycomb-mediated repression at keratinocyte differentiation genes is unaltered upon *ACTL6A* over-expression. Related to Figure 5.**
(A) Genome browser tracks of H3K27me3 CUT&RUN and RNA-seq profiles at keratinocyte-differentiation gene *KRT1* and its nearby genes (left), and *LOR* (right) in primary human keratinocytes transduced by lentivirus overexpressing *ACTL6A* and the vector control. Also shown were H3K27me3 and H3K4me3 ChIP-seq tracks in keratinocytes from ENCODE datasets. (B) Expression levels (transcripts per million, TPM) of keratinocyte differentiation genes and progenitor markers between *ACTL6A*-overexpressed and vector-control keratinocytes. (C) Genome browser tracks as in (A) at the *HOXB* locus.
